# Bridging the gap between single-cell migration and collective dynamics

**DOI:** 10.1101/548677

**Authors:** Florian Thüroff, Andriy Goychuk, Matthias Reiter, Erwin Frey

**Affiliations:** Arnold Sommerfeld Center for Theoretical Physics and Center for NanoScience, Department of Physics, Ludwig-Maximilians-Universität München, Theresienstr. 37, D-80333 Munich, Germany

## Abstract

A wealth of experimental data relating to the emergence of collective cell migration as one proceeds from the behavioral dynamics of small cohorts of cells to the coordinated migratory response of cells in extended tissues is now available. Integrating these findings into a mechanistic picture of cell migration that is applicable across such a broad range of system sizes constitutes a crucial step towards a better understanding of the basic factors that determine the emergence of collective cell motion. Here we present a cellular-automaton-based modeling framework, which focuses on the integration of high-level cell functions and their concerted effect on cellular migration patterns. In particular, we adopt a top-down approach to incorporate a coarse-grained description of cell polarity and its response to mechanical cues, and address the impact of cell adhesion on collective migration in cell groups. We demonstrate that the model faithfully reproduces typical cell shapes and movements down to the level of single cells, yet is computationally efficient enough to allow for the simulation of (currently) up to 𝒪(10^4^) cells. To develop a mechanistic picture that illuminates the relationship between cell functions and collective migration, we present a detailed study of small groups of cells in confined circular geometries, and discuss the emerging patterns of collective motion in terms of specific cellular properties. Finally, we apply our computational model at the level of extended tissues, and investigate stress and velocity distributions, as well as front morphologies, in expanding cellular sheets.

---

Cell movements range from uncoordinated ruffling of cell boundaries to the migration of single cells [1] to the collective motions of cohesive cell groups [2]. Single-cell migration enables cells to move towards and between tissue compartments – a process that plays an important role in the inflammation-induced migration of leukocytes [3]. One can distinguish between amoeboid and mesenchymal migration, which are characterized by widely different cell morphologies and adhesive interactions with their respective environments [4, 5]. Cells may also form cohesive clusters and mobilize as a collective [6–11]. This last mode of cell migration is known to drive tissue remodelling during embryonic morphogenesis [12] and wound repair [13].

Despite this broad diversity of migration modes, there appears to be a general consensus that all require (to varying degrees) the following factors: (i) Cell polarization, cytoskeletal (re)organization, and force generation driven by the interplay between actin polymerization and contraction of acto-myosin networks. (ii) Cell-cell cohesion and coupling mediated by adherens-junction proteins which are coupled to the cytoskeleton. (iii) Guidance by chemical and physical signals. The basic functionalities implemented by these different factors confer on cells the ability to generate forces, adhere (differentially) to each other and to a substrate, and respond to mechanical and chemical signals. However, a fully mechanistic understanding of how these basic functionalities are integrated into single-cell migration and coordinated multicellular movement is still lacking.

Here, we present a computational model which enables us to study cell migration at various scales, and thus provides an integrative perspective on the basic cell functions that enable the emergence of collective cell migration. While a variety of very successful modeling approaches has been used to describe single-cell dynamics [14–23] or the movements of extended tissues [24–31], these models are hard to reconcile with each other. Models that focus on single cells are typically difficult to extend to larger cell numbers, largely due to their computational complexity. On the other hand, approaches which are designed to capture the dynamics at the scale of entire tissues generally adopt a rather coarse-grained point of view, and are therefore difficult to transfer to single cells or small cell cohorts. At present there are two partly competing and partly complementary approaches to bridge the gap between single-cell migration and collective dynamics, namely phase-field models [17, 18, 32–35], and cellular Potts models (CPMs) [25, 26, 36–41] first introduced by Graner and Glazier [42].

Building on and generalizing the CPM [42], we present a cellular automaton model that is designed to capture essential cellular features even in the context of the migration of single cells and of small sets of cells. At the same time, it is computationally efficient for simulations with very large cell numbers [currently up to 𝒪(10^4^) cells], thus permitting investigations of collective dynamics at the scale of tissues. Our model reproduces the most pertinent features of cell migration even in the limiting case of solitary cells, and is compatible with a wealth of experimental evidence derived from both small cell groups and larger collectives made up of several thousand cells. Specifically, by studying the characteristics of single-cell trajectories and of small cell groups confined to circular territories, we demonstrate that persistency of movements is significantly affected by cell stiffness and cell polarizability. Moreover, we investigate the dynamics of tissues in the context of a typical wound-healing assay [6, 13, 43], and show that the model exhibits the *X*-shaped traction force patterns observed experimentally [43], a feature which we attribute to the coupling between cell-sheet expansion and cell-density-induced growth inhibition.

## I. COMPUTATIONAL MODEL

We consider a cell *α* as a set of simply connected grid sites 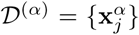 on a two-dimensional (2D) lattice (Fig. 1). Cell motion and cell shape changes correspond to the annexation or relinquishment of grid sites, and, as for actual cells, involve protrusions (retractions) of cell boundaries [44, 45]. In the computational model, these processes are implemented as elementary protrusion and retraction events, 𝒯_pro_ and 𝒯_ret_, corresponding to the increase and decrease, respectively, of the number of lattice elements within 𝒟^(*α*)^ by takeover (surrender) of individual grid sites at the cell’s boundary ℬ^(*α*)^. The outcome of each takeover attempt is determined on the basis of its associated cost or payoff in some configuration energy assigned by a Monte Carlo scheme. The key mechanical structures driving all these processes include branched actin networks and actin bundles, acto-myosin networks, adherens junctions, and focal adhesions. In our formalism, we model the effects of these structures in a coarse-grained manner as follows.

**FIG. 1.**
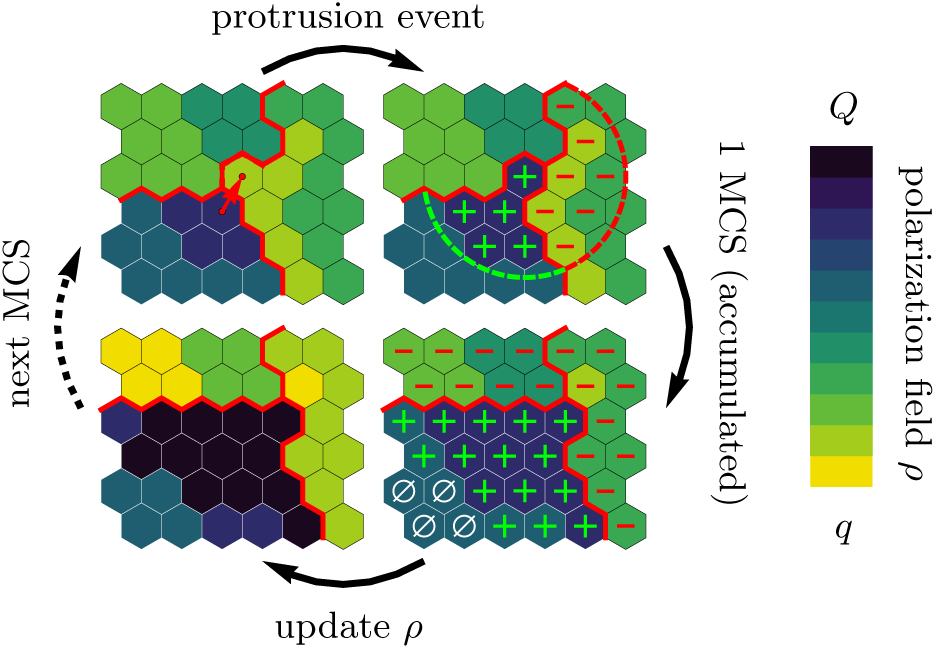
Illustration of the computational model with the pertinent simulation steps. The scheme depicts three cells (bounded by the red lines), each occupying a contiguous set of grid sites (hexagons). *Top left:* The upper right corner of the lower left cell (source cell) initiates a protrusion event against a neighboring element in the cell to its right (target cell), as indicated by the arrow, in an attempt to displace it. The success of each such attempted elementary event depends on the balance between contractile forces (ℋ _cont_), cytoskeletal forces (ℋ_cyto_), and cell adhesion (ℋ_adh_). *Top right:* If the protrusion event is successful, the levels of regulatory factors are increased (decreased) in integer steps at all lattice sites inside the source (target) cell, that lie within a radius *R* of the accepted protrusion event (as indicated by the plus and minus signs). *Bottom right:* During the course of one MCS, different levels of regulatory factors accumulate locally within each cell, with positive levels of regulatory factors (green plus signs) promoting a build-up of cytoskeletal structures, negative levels of regulatory factors (red minus signs) causing degradation of cytoskeletal structures, and neutral levels of regulatory factors (white zero signs) causing relaxation towards a resting state, as indicated in the *lower left image*. The color code indicates local levels of cytoskeletal structures, *ρ*.

As in the CPM, we assume that deformations of a cell’s membrane and cortex are constrained by the elastic energy

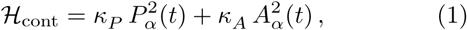

where *κ*_*P*_ and *κ*_*A*_ are cell-type-specific stiffness parameters for the perimeter *P*_*α*_(*t*) and the area *A*_*α*_(*t*) of a cell *α* at time *t*, respectively. The ensuing contractile forces are counteracted by outward pushing forces generated by assembling and disassembling cytoskeletal structures [14, 44]. To model these dynamic processes we generalize the CPM by introducing a time-dependent and spatially resolved internal concentration field for each cell: *ρ*_*α*_(**x**_*n*_, *t*), which is intended to represent the density of force-generating cytoskeletal structures. We assume that for an elementary protrusion event the energy change is given by the difference in this density between target site **x**_*t*_ and source site **x**_*s*_,

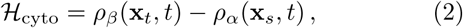

where the amplitude of *ρ*_*α*_ encodes for the energy scale; an analogous expression holds for elementary retraction events.

Assembly and disassembly of cytoskeletal structures are controlled by a myriad of accessory proteins [1, 45]. Since there are several biological factors which limit the local density of actin filaments, we introduce cell-type-specific bounds for the cytoskeletal field: *q* ≤ *ρ*(**x**_*n*_, *t*) ≤*Q*. While the upper bound *Q* mainly reflects the limited availability of proteins, the lower bound *q* serves to prevent cells from collapsing. Moreover, cytoskeletal structures are known to respond to external mechanical stimuli through feedback mechanisms involving regulatory cytoskeletal proteins[15, 16]. In our computational model, we greatly simplify these complex processes by subsuming them into a single integer variable *F* (**x**_*n*_, *t*) which we will refer to as “*regulatory factors*”. If a protrusion or retraction event has been accepted at some source site, then, in a process that could be called a *mechanotransduction* mechanism, *F* (**x**_*n*_, *t*) is within some radius *R* accordingly altered up or down for the protruding and the retracting cell, respectively. These regulatory factors in turn modulate the assembly and disassembly of cytoskeletal structures. Specifically, we assume that positive levels, *F* (**x**_*n*_, *t*) > 0, promote assembly

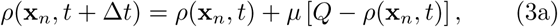

while negative values, *F* (**x**_*n*_, *t*) < 0, favor disassembly

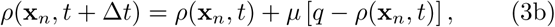

and neutral values, *F* (**x**_*n*_, *t*) = 0, favor relaxation towards a resting state

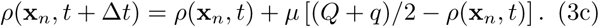

The parameter *µ* signifies the rate at which cytoskeletal structures respond to the regulatory factors.

In addition to internal remodeling of the cytoskeleton, adhesion of cells to neighboring cells and to the substrate plays a key role in explaining migratory phenotypes [2, 14]. From a mechanical point of view, the implications of cell adhesion are two-fold. First, cell adhesion supports growth of cell-cell and cell-matrix contacts and may thus be described in terms of effective surface energies. Secondly, once formed, adhesive bonds anchor the cell to the substrate and to neighboring cells. During cell migration, these anchoring points must continuously be broken up and reassembled [46, 47] and, hence, provide a constant source of dissipation. In our computational model, the dissipative nature of cell-cell adhesions is accounted for as follows: while the formation of new cell-cell adhesions is favored by an energy benefit *B*, the rupture of such cell-cell adhesions is accompanied by an energy cost *B* + Δ*B* > *B*, where Δ*B* represents the dissipative nature of cell-cell adhesions.

To model cell-substrate adhesion, we introduce a second scalar field *φ*(**x**) whose value is taken to reflect the density of substrate sites at which focal adhesions between cell and substrate can be formed. By allowing *φ*(**x**) to take negative values, we can, moreover, define lattice areas with cell-repelling properties, thus providing a natural means of implementing arbitrary substrate micropatterns. Analogously to cell-cell contacts, we account for the dissipative nature of cell-substrate adhesions by associating the breaking of such contacts with an additional energy cost *D*.

In the description so far, the cells are arrested in the cell cycle (mitostatic). To investigate the effect of cell proliferation on tissue dynamics, we introduce a simplified three-state model of cell division. Cells start off in a quiescent state, in which their properties remain constant over time. The cell sizes fluctuate around an average value determined by the cell properties and the local tissue pressure. Upon exceeding a threshold size (*A*_0_) due to size fluctuations, cells leave the quiescent state and enter a growth state. The duration of the quiescent state is thus a random variable, whose average value depends on the tissue pressure, and lower pressure leads to a shorter quiescent state. During its subsequent deterministic growth state of duration *T*_*g*_, the cell doubles all of its cellular material and thus doubles in size. We model this growth as a gradual decrease in the effective cell contractility (*κ*_*A*_ and *κ*_*P*_). As there is no a priori reason to assume that the cell’s migratory behavior should depend on its size, we constrain the parameters accordingly; this is described in detail in the Supplemental Material [48]. After having grown, the cells switch to the deterministic division state of duration *T*_*d*_. Here, each cells utilizes its cytoskeleton for the separation of the cellular material instead of migration, thus reducing cell polarizability to zero: Δ*Q* → 0. At the end of the division state, each dividing cell splits into two identical daughter cells. The daughter cells’ properties and parameters are identical to the mother cell’s initial values in the quiescent state. For a detailed and more technical description we refer the interested reader to the Supplemental Material [48].

## II. RESULTS

### A. Single-cell migration

To study the migration of single cells, we used a large computational grid with 9·10^4^ sites and periodic boundary conditions. The average cytoskeletal density was fixed at (*q* + *Q*)/2 = 225. We then investigated the impact of varying cell perimeter stiffness *κ*_*P*_ and levels of maximum cell polarity Δ*Q* ≡ *Q* – *q* on the cell’s migratory patterns. To assess the statistics of the cell trajectories, we recorded the cell’s orientation 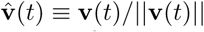 (**v**: cell velocity) and (geometrical) center of mass position **R**(*t*) during a total simulation time of *T* = 10^4^ Monte-Carlo steps (MCS). For each set of parameters, we performed 100 independent simulations, from which we computed the mean squared displacement, MSD(*τ*) ≡ ⟨ [**R**(*t* + *τ)* − **R**(*t*)]^2^⟩, and the normalized velocity autocorrelation function, 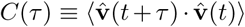, where ⟨ *…* ⟩denotes an average with respect to simulation time *t* as well as over all 100 independent simulations.

These computer simulations show that the statistics of the migratory patterns is well described by a *persistent random walk model* [49, 50] with its two hallmarks: a mean square displacement that exhibits a crossover from ballistic to diffusive motion [Fig. 2], and on sufficiently long time scales an exponential decay of the velocity autocorrelation function 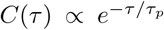 [inset of Fig. 2].

**FIG. 2.**
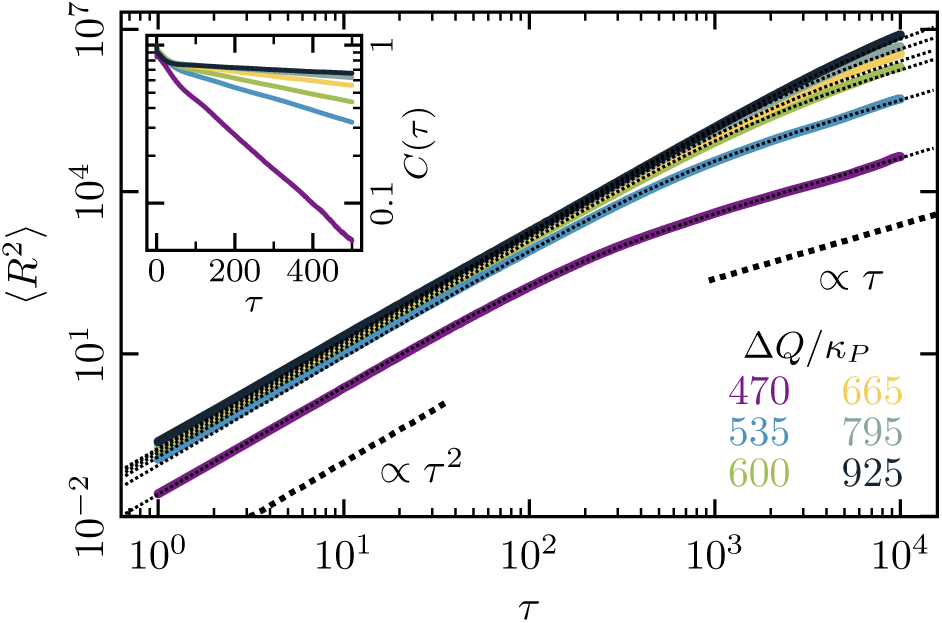
Mean-squared displacement (MSD) for single-cell movements at different maximum cell polarity Δ*Q* (stiffness parameters *κ*_*P*_ = 0.060, *κ*_*A*_ = 0.18; average cytoskeletal density (*Q*+*q*)/2 = 225; signaling radius *R* = 5; cell-substrate dissipation *D* = 0; cell-substrate adhesion penalty *φ* = 0). Single cells perform a persistent random walk, i.e. they move ballistically (MSD ∝ *τ* ^2^) for *τ* ≪ *τ* _*p*_, and diffusively (MSD ∝ *τ)* for *τ* ≫*τ*_*p*_. *Inset:* Normalized velocity auto-correlation function for the same parameters as in the main figure.

The persistence time *τ*_*p*_ has a characteristic dependence on the maximum cell polarity Δ*Q*. There is a threshold value for Δ*Q* below which cells remain immobile. Above this threshold, the persistence time *τ*_*p*_ shows a marked increase with Δ*Q* [Fig. 3A]: cells with larger Δ*Q* exhibit extended episodes of ballistic motion. Likewise, persistence times increase as the cell membrane and cortex become more compliant (as the value for the perimeter stiffness *κ*_*P*_ is reduced). Interestingly, our simulations also show that there is a strong correlation between cell shape and the persistence time of the cell’s trajectory [Fig. 3B]. While highly persistent trajectories are observed for cells with ‘crescent’ shapes, more erratic cell motion is typically found for cells with more rounded outlines. In other words, our computational model predicts that cells which are able to polarize their cytoskeletal structures more strongly will adopt crescent shapes and show a higher degree of persistent cell motion. It would be interesting to test these predictions using phenotypic variations in cell shapes like those reported in experiments with keratocytes [51].

**FIG. 3.**
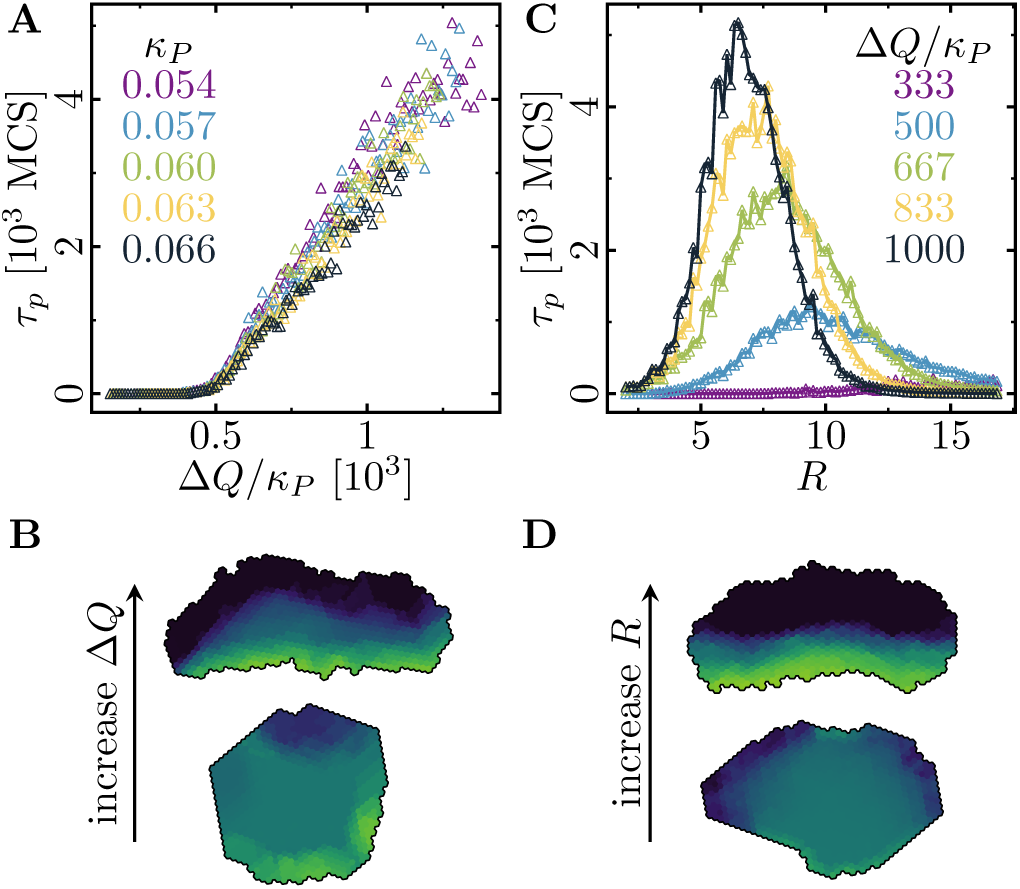
Cell shape and persistence of migration as a function of the polarization parameters. **(A)** Persistence times of persistent random walks by single cells are plotted as a function of maximum cell polarity Δ*Q*, and perimeter stiffness *κ*_*P*_ (area stiffness *κ*_*A*_ = 0.18; average cytoskeletal density (*Q* + *q*)/2 = 225; signaling radius *R* = 5; cytoskeletal update rate *µ* = 0.1). The persistence time of the random walk increases with increasing cytoskeletal polarity and decreasing perimeter elasticity. **(B)** Cytoskeletal polarity also controls cell shapes, with crescent cell shapes (long persistence times) being observed at large cytoskeletal polartities, and elongated cell shapes (short persistence times) at small cytoskeletal polarities. **(C)** Persistence times of persistent random walks by single cells as a function of the cell’s signaling radius at different values for the cytoskeletal polarity (stiffness parameters *κ*_*P*_ = 0.060, *κ*_*A*_ = 0.18; average cytoskeletal density (*Q* + *q*)/2 = 225; cytoskeletal update rate *µ* = 0.1). **(D)** The signaling radius critically determines the synchronicity of internal cytoskeletal remodeling processes. Small signaling radii frequently lead to transient formation of mutually independent lamellipodia at different positions around the cell perimeter, thereby interrupting persistent motion (reducing persistence times). Large signaling radii lead to structurally stable front-rear polarization profiles across the entire cell body (long persistence times). **(B)**, **(D)** Color code: cell polarization; cf. color bar in Fig. 1.

Moreover, we investigated the influence of different signaling radii *R* on the persistence of single-cell trajectories. Since *R* controls the spatial organization of lamellipodium formation, its value should strongly affect the statistics of a cell’s trajectory [Fig. 3C]. Indeed, at small values of *R*, we observe that the spatial coherence of cytoskeletal rearrangements is low, which frequently results in the disruption of ballistic motion due to the formation of independent lamellipodia in spatially separate sectors of the cell boundary [Fig. 3D, upper snapshot]. In contrast, at larger values of *R*, we find that spatial coherence is restored, and the formation of one extended lamellipodium across the cell’s leading edge maintains a distinct front-rear axis of cell polarity [Fig. 3D, lower snapshot].

### B. Circular micropatterns

To assess the transition to collective cell motion, we next studied the dynamics of small cell groups confined to circular micropatterns [8, 9, 38, 52]. We implemented these structures in silico by setting *φ*(**x**) = 0 inside a radius *r*_0_ and *φ*(**x**) → -∞ outside. During each simulation run, the number of cells was also kept constant by deactivating cell division. We previously employed this setup to compare our numerical results with actual experimetal measurements, and found very good agreement [38]. Here, we generalize these studies and present a detailed analysis of the statistical properties of the collective dynamics of cell groups in terms of the key parameters of the computational model.

When adhesive groups of two or more motile cells are confined on a circular island, they arrange themselves in a state of spontaneous collective migration, which manifests itself in the form of coordinated and highly persistent cell rotations about the island’s midpoint **x**_0_ [8, 9, 38, 52]. The statistics of these states of rotational motion provide insight into the influence of cellular properties on the group’s ability to coordinate cell movements. To quantify cell rotations in a system of *K* cells, we recorded the angular distance *φ*_*α*_(*t*) covered by each cell *α* over time and determined the average angular trajectory *Φ*(*t*) ≡ *K*^*-*1^ Σ_*α*_ *φ*_*α*_(*t*) traversed by each cell. The resulting random variables for the overall angular velocity of the cell assembly, 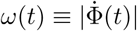, and the average cell perimeter *P* (*t*) ≡ *K*^*-*1^Σ_*α*_ *P*_*α*_(*t*) were then used to characterize the statistics of collective cell rotation. For each specific choice of simulation parameters, we monitored *ω*(*t*) and *P* (*t*) for a set of 10 statistically independent systems, each of which was observed over *T* = 10^4^ MCS. From these data, we then computed the mean overall rotation speed ⟨*ω*⟩, its variance 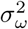, and the variance of the cell perimeter, 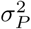.

Figure 4 illustrates the characteristic properties of collective cell rotations in systems containing *K* = 4 cells and endowed with varying maximum cell polarity Δ*Q* and varying cell contractility *κ*_*P*_. Interestingly, the statistical measures shown in Fig. 4A do not separately depend on cell contractility and maximum cell polarity, but depend only on their ratio Δ*Q/m*, which we will henceforth refer to as *“specific polarity”*. Overall, we observed that upon increasing the specific polarity there is a marked transition from a quiescent state to a state where the cells are collectively moving. Below a threshold value for the specific polarity [Δ*Q/m*≈1300 in Fig. 4], the rotation speed 〈*ω*〉 [purple curves in Fig. 4A] equals zero and the cells are immobile. In this regime, which we term the *“stagnation phase”*, or 𝒮-phase, cytoskeletal forces are too weak to initiate coherent cell rotation, and the system’s dynamics is dominated by relatively strong contractile forces, which tend to arrest the system in a “low energy” configuration. Beyond this threshold, we identified three distinct phases of collective cell rotation. In the ℛ_1_-phase, we observed a steep increase in the average rotation speed and a local maximum in the fluctuations of both cell shape and rotation speed; cf. green (*σ*_*P*_) and blue (*σ*_*ω*_) curves in Fig. 4A. Now, cytoskeletal forces are sufficiently large to establish actual membrane protrusions against the contractile forces, and cells begin to rotate [Fig. 4B,C]. However, the contractile forces still dominate, such that cellular interfaces tend to straighten out and lamellipodium formation is sustained only over finite lifetimes. Thus, due to the dominance of contractile forces, the systems frequently experience transient episodes of stagnation and repeatedly change direction [Fig. 4B].

**FIG. 4.**
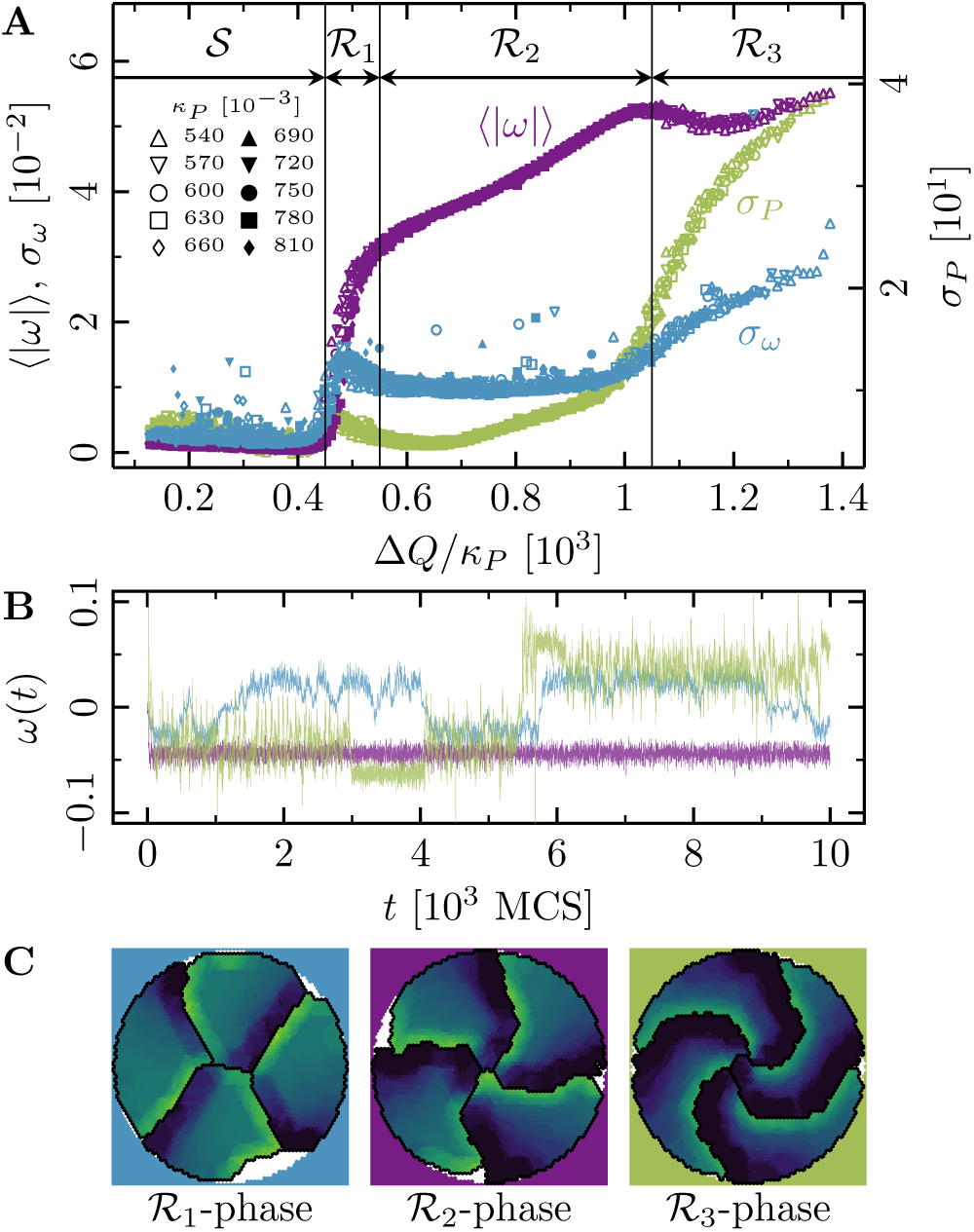
Phases of collective motion. (4-cell systems, confinement radius *r*_0_ = 30.6; area stiffness *κ*_*A*_ = 0.18; average cytoskeletal density (*Q* + *q*)*/*2 = 225; signaling radius *R* = 5; cytoskeletal update rate *μ* = 0.1; cell-cell adhesion *B* = 0; cell-cell dissipation Δ*B* = 7; cell-substrate dissipation *D* = 0; cell-substrate adhesion penalty *φ* = 0 (*r < r*_0_), *φ* →*-∞* (*r > r*0)). **(A)** Characteristic observables of collective cell rotation at different values of the cell perimeter stiffness parameter *κ*_*P*_ : mean (⟨*ω*⟩)and standard deviation (*σ*_*ω*_) of angular velocity of cell motion, and the standard deviation of the cell shape variability (*σ*_*P*_). The statistics of collective cell motion depends only on the ratio of maximum cell polarity, Δ*Q*, to cell contractility, *κ*_*P*_ (specific polarity). **(B)** Representative angular trajectories and **(C)** cell shapes (color code represents cell polarization; cf. Fig. 1) for the different parameter regimes as described in the main text.

At intermediate values of the specific polarity (ℛ_2_-phase), the cellular systems reach a regime of enduring rotational motion, where ⟨*ω*⟩ varies linearly with the specific local polarity, and where *σ*_*P*_ and *σ*_*ω*_ exhibit a rather broad minimum [Fig. 4A]. In this regime, a range of “optimal ratios” of cytoskeletal to contractile forces sustains stable cell shapes, and sets the stage for the formation of extended lamellipodia and the establishment of permanent front-rear polarizations of cells. As a result, persistence times become very large, rendering cellular rotations strictly unidirectional within the observed time window [Fig. 4B]. Finally, at large values of the specific polarity (ℛ_3_-phase), the system’s dynamics is dominated by cytoskeletal forces and the rotational speed ⟨*ω*⟩ saturates at some maximal value. Due to the relatively small contractile forces, cell shapes tend to become unstable, as reflected in the growing variance of the cell perimeter *σ*_*P*_ [green curve in Fig. 4A]. These instabilities in cell shape frequently lead to a loss of persistence in the rotational motion of the cells [growing *σ*_*ω*_; blue curve in Fig. 4A].

### C. Tissue-level dynamics

As an application of our computational model at the tissue level, we considered a setup in which an epithelial cell sheet expands into free space. As in recent experimental studies [6, 13, 27, 43], we confined cells laterally between two fixed boundaries, within which they proliferated until they reached confluence; in the *y*-direction we imposed periodic boundary conditions. Then we removed the boundaries and studied how the cell sheet expands. In order to quantify tissue expansion, we monitored cell density and velocity, as well as the mechanical stresses driving the expansion process. Figure 5 shows our results for two representative parameter regimes that highlight the difference between a dynamics dominated by cell motility in the absence of cell proliferation, and a contrasting regime where low-motility cells grow and divide depending on the local cell density.

**FIG. 5.**
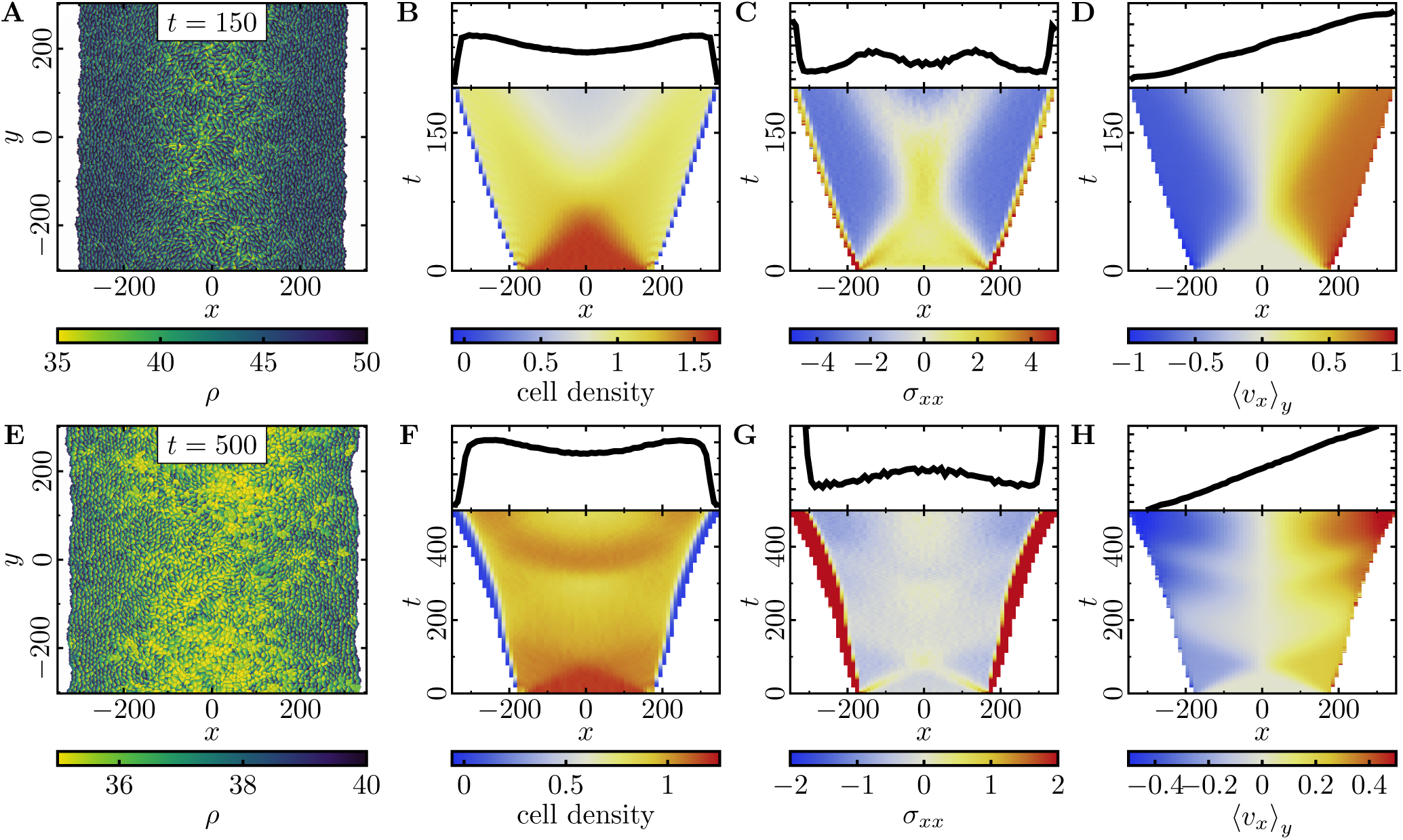
**Expansion of a confluent epithelial cell sheet** after removal of boundaries positioned at *x* = *±*175 for two different parameter settings. **(A-D)** Tissue expansion for a migration-dominated setup without explicit cell growth and mitosis. (3300-cell system; stiffness parameters *κ*_*P*_ = 0.12, *κ*_*A*_ = 0.18; average cytoskeletal density (*Q* + *q*)*/*2 = 35; maximum cell polarity Δ*Q* = 30; signaling radius *R* = 2; cytoskeletal update rate *μ* = 0.1; cell-cell adhesion *B* = 7; cell-cell dissipation Δ*B* = 0; cell-substrate dissipation *D* = 0; cell-substrate adhesion penalty *φ* = 0). **(E-H)** Tissue expansion at low density and cell polarizability for a cell sheet comprised of dividing cells. (Initially a 2500-cell system; stiffness parameters *κ*_*P*_ = 0.12, *κ*_*A*_ = 0.18; average cytoskeletal density (*Q* + *q*)*/*2 = 35; maximum cell polarity Δ*Q* = 10; signaling radius *R* = 2; cytoskeletal update rate *μ* = 0.1; cell-cell adhesion *B* = 7; cell-cell dissipation Δ*B* = 0; cell-substrate dissipation *D* = 0; cell-substrate adhesion penalty *φ* = 0; growth time *T*_*g*_ = 180; division time *T*_*d*_ = 20; size threshold for cell growth *A*_T_ = 1 *A*_ref_, where *A*_ref_ is the size of a solitary cell in equilibrium). **(A, E)** Snapshots of the density of cytoskeletal structures *ρ*. **(B, F)** Kymographs showing the cell density averaged over the *y*-direction for and (*top*) final snapshots of the cell density profiles. **(C, G)** Kymographs showing the component *σ*_*xx*_ of the stress tensor averaged over the *y*-direction and (*top*) final snapshots of the stress profiles. **(D, H)** Kymographs showing the component *v*_*x*_ of the cell velocities averaged over the *y*-direction and (*top*) final snapshot of the velocity profiles.

We first investigated how a densely packed pre-grown tissue of mitostatic cells with high polarizability (large Δ*Q*) expands into cell-free space upon removal of the confining boundaries at the tissue’s lateral edges [Fig. 5A]. As the cells migrate into the cell-free space, we observe a strongly (spatially) heterogeneous decrease in the initially high and uniform cell density and mechanical pressure in the expanding monolayer [Fig. 5B, C]. This is quite distinct from the behavior of a homogeneous and ideally elastic thin sheet, which would simply show a homogeneous relaxation in density as it relaxes towards its rest state. Moreover, cell polarization and the ensuing active cell migration lead to inhomogeneously distributed traction stresses in the monolayer. After initial expansion of the monolayer, facilitated by high mechanical pressure, the cells at the monolayer edge begin to polarize outwards, which enhances outward front migration. These actively propagating cells exert traction the trailing cells, and thereby yield a trailing region with negative stress [Fig. 5C]. Taken together, this gives rise to a characteristic X-shaped pattern in the kymograph of the total mechanical stresses ⟨*σ*_*xx*_⟩ _*y*_ [Fig. 5C]. This profile closely resembles the first period of mechanical waves observed experimentally [43]. It illustrates how stress is transferred towards the center of the monolayer when cells are highly motile and collectively contribute to tissue expansion. At the end of the simulated time window, the cell density exhibited a minimum in the center of the sheet [Fig. 5C]. This is due to stretching of the central group of cells caused by the equally strong traction forces exerted by their migrating neighbors on both sides. Finally, the simulations also showed that outward cell velocities increased approximately linearly with distance from the center, confirming that the entire cell sheet contributes to the expansion in this configuration.

To explore the possible range of tissue dynamics and expansion, we also investigated a qualitatively different parameter regime where cells are less densely packed and can also polarize less due to a narrower range of polarizability [Fig. 5E, F]. Here, the expansion of the monolayer is mainly driven by cell division, and cells keep dividing until they reach a homeostatic cell density. Even though cells should typically exceed the threshold size and hence enter the growth phase at different times, we observe that the cell sheet exhibits periodic ‘bursts’ of growth coinciding with the total duration of a complete cell cycle (200 MCS) and alternating with cell migration [Fig. 5H]. These periodic ‘bursts’ can be explained as follows. Initially, the slightly compressed monolayer expands to relieve mechanical pressure. Due to this initial motion, the cells at the monolayer edge begin to polarize outwards. As in the previous case, where cell proliferation is absent [Fig. 5A-D], the polarized cells enhance outward front migration and stretch the cells in the bulk of the cell sheet. For the same reasons as before, we observe a typical X-shaped stress pattern in the kymograph [Fig. 5G], albeit less pronounced due to the lower polarizability of the cells [cf. Fig. 5C]. Because a broad region of cells in the monolayer bulk is stretched by the actively migrating cell fronts, these cells exceed the threshold size and begin growing approximately in phase. Once the mechanical pressure of the cell sheet is relieved, it will stop expanding [Fig. 5H]. However, cell growth and division once more lead to an increase in mechanical pressure (and cell density) in the monolayer. This cycle of migration-dominated monolayer expansion and cell-density-dependent cell growth and division results in a periodic reoccurence of the X-shaped stress pattern [Fig. 5G], closely resembling the pattern observed in experiments [43].

On a sidenote, the mentioned synchronization of the cell division and cell migration phases by the deterministic portion of the cell cycle can be counteracted by introducing additional stochastic terms in the transition between the different phases of the cell cycle.

## III. CONCLUSION

In this work, we have proposed a generalization of the cellular Potts model [42]. The model implements a coarse-grained routine that captures the salient features of cytoskeletal remodeling processes on subcellular scales, while being computationally tractable enough to allow for the simulation of entire tissues containing up to 𝒪(10^4^) cells. We have used the model to study the transition from single-cell to cohort cell migration in terms of the interplay between the pertinent cellular functions. Specifically, we have demonstrated that our model consistently reproduces the dynamics and morphology of motile cells even down to the level of solitary cells. Our studies also reveal that cytoskeletal forces (relative to cell contractility), as well as the spatial organization of the cells’ lamellipodia, significantly affect the persistency of cellular trajectories, both in the context of single-cell motion and in cohesive cell groups restricted to circular micropatterns. On larger scales, our simulation results suggest that the dynamics of expanding tissues strongly depends on the specific properties of the constituent cells. If monolayer expansion is driven by active cell migration throughout the tissue, then the cell sheet exhibits typical traction-force patterns and an X-shape in the kymograph. Additionally, a cell-density-dependent cell growth leads to a periodic recurrence of these traction-force patterns in a cycle of migration-dominated expansion and ‘burst’-like cell proliferation.

Taken together, our results further highlight the intricacies of collective cell migration, which involves a multitude of intra- and inter-cellular signaling mechanisms operating at different scales in length and time. Establishing a comprehensive picture that incorporates and elucidates the mechanistic basis of these phenomena remains a pressing and challenging task. The multiscale modeling approach proposed here provides a direct link between subcellular processes and macroscopic dynamic observables, and might thus offer a viable route towards this goal.

## ACKNOWLEDGMENTS

A.G. acknowledges support by a DFG fellowship through the Graduate School of Quantitative Biosciences Munich (QBM). E.F. acknowledges support by the German Excellence Initiative via the program ‘NanoSystems Initiative Munich’ (NIM) and by the Deutsche Forschungsgemeinschaft (DFG) via Collaborative Research Center (SFB) 1032 (project B02). We thank Felix Kempf, Felix Segerer, Sophia Schaffer and Joachim Rädler for stimulating discussions.

