## Supplemental Text for "Bridging the gap between single-cell migration and collective dynamics"

### CONTENTS

S.I. Computational model 1
A. Computational grid 1
B. Representation of biological cells 2
C. Model dynamics 2
1. Protrusion and retraction of cells 2
2. Monte-Carlo scheme 2
D. Implementation of cellular traits 3
1. Cell contractility 3
2. Cytoskeletal structures and focal adhesion 4
3. Cell adhesion 6
4. Rupture of cell contacts 7
5. Dissipation in substrate adhesion 8
E. Cell domain update routine 9
F. Cell proliferation and mitosis 9
G. Numerical computation of stress in a tissue 10
H. Numerical computation of the cell shape 11

S.II. Parameter screening in silico 12
A. Single cell size 12
B. Single cell shape and dynamics 13
C. Cells in circular confinement 15
D. Velocity and roughness of spreading tissue 16

References 19

### S.I. COMPUTATIONAL MODEL

In this section, we describe in detail the implementation of our computational model, which has been outlined briefly in the main text. While the biological rationale behind our modeling approach has been discussed in the main text, our focus here is on the technical aspects and the details of the numerical implementation. To facilitate subsequent discussions on implementation details, we start by introducing some model-specific terminology which will be used throughout this section to illustrate the mechanics of our model.

#### A. Computational grid

The basic data structure, our computational model resorts to, is referred to as the *grid*; see Fig. S1. The grid itself is implemented as a regular, space-filling lattice with lattice vectors $\{\mathbf{x}_i\}_{i=1,\dots,N}$. Each lattice vector $\mathbf{x}_i$ is understood to represent its associated Voronoi cell which will be referred to as *grid site*. To be specific, we consider triangular tilings $\{\mathbf{x}_i\}_{i=1,\dots,N}$, such that each grid site is a hexagon, which is surrounded by 6 nearest-neighbor sites that define the neighborhood $\mathcal{N}_k$ of $\mathbf{x}_k$:

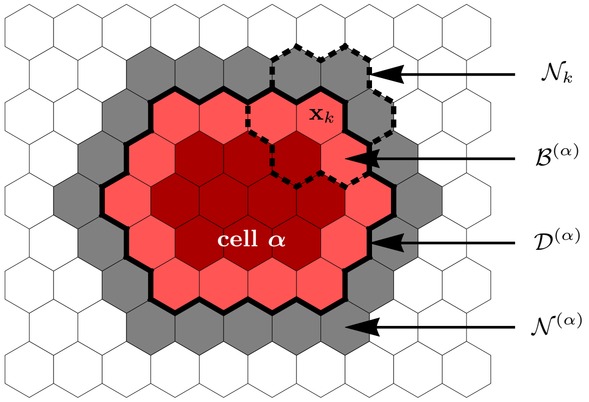


FIG. S1. Illustration of the various sets defining a cell and its environment. Grid sites occupied by cell $\alpha$, i.e. its domain $\mathcal{D}^{(\alpha)}$, are indicated in red colors. The cell's membrane sites, $\mathcal{B}^{(\alpha)}$, are indicated by the lighter red color, the cell's immediate neighborhood, $\mathcal{N}^{(\alpha)}$, is indicated in gray color. Elementary events involving cell $\alpha$ always involve one grid site in $\mathcal{B}^{(\alpha)}$ and one grid site in $\mathcal{N}^{(\alpha)}$. For the hexagonal lattices used in this work, each grid site $\mathbf{x}_k$ is surrounded by 6 nearest neighbors which we collectively denote by $\mathcal{N}_k$.

$$\mathcal{N}_k = \Big\{\mathbf{x}_j \,\Big|\, \mathbf{x}_j \text{ is nearest neighbor of } \mathbf{x}_k\Big\}. \tag{S1}$$

Overall, the grid represents our general notion of (discretized) space, and each grid site holds information specific to cells as well as to environmental factors. In what follows, distances on this spatial grid will be measured in units of the distance between the midpoints of neighboring lattice sites, i.e.

$$||\mathbf{x}_k - \mathbf{x}_j|| = 1 \Leftrightarrow j \in \mathcal{N}_k. \tag{S2}$$

This then implies for the side length $\ell$ and the two-dimensional volume (area) $a$ of each hexagonal grid site: $\ell = 1/\sqrt{3}$ and $a = 3\sqrt{3}\,\ell^2/2$.

---

* These two authors contributed equally.

#### B. Representation of biological cells

In the spirit of the cellular Potts model [1, 2], each cell is represented by a simply connected set of lattice sites

$$\mathcal{D}^{(\alpha)} = \Big\{ \mathbf{x}_k \Big| \, c(\mathbf{x}_k) = \alpha \Big\}, \tag{S3}$$

where the indicator function $c(\mathbf{x}_k)$ gives the index of the cell occupying $\mathbf{x}_k$. Here and in the following, we use latin indices to reference lattice sites, and greek indices to reference cells. The set $\mathcal{D}^{(\alpha)}$ used to represent the spatial extension of cell $\alpha$, will be referred to as the *domain* of cell $\alpha$. In our model, each grid site, $\mathbf{x}_k$, can be occupied by at most one cell (i.e. we do not allow for overlapping cell domains). The absence of cells at $\mathbf{x}_k$ is numerically implemented by negative values of the indicator function, $c(\mathbf{x}_k) < 0$. Following this terminology, the area and the perimeter of cell $\alpha$ is given by:

$$A_\alpha = a \sum_{k=1}^{N} \delta_{\alpha, c(\mathbf{x}_k)} = \frac{3\sqrt{3}}{2} \ell^2 \sum_{k=1}^{N} \delta_{\alpha, c(\mathbf{x}_k)}, \tag{S4a}$$

$$P_\alpha = \ell \sum_{k=1}^{N} \sum_{\mathbf{x}_l \in \mathcal{N}_k} \delta_{\alpha, c(\mathbf{x}_k)} \big(1 - \delta_{\alpha, c(\mathbf{x}_l)}\big). \tag{S4b}$$

#### C. Model dynamics

##### 1. *Protrusion and retraction of cells*

Biological cells are highly dynamic entities which constantly change shape and move around in space. To reflect this dynamic behavior computationally, the domain $\mathcal{D}^{(\alpha)}$ of cell $\alpha$ changes over time. The evolution of cell shape and position, as represented by $\mathcal{D}^{(\alpha)}$, proceeds via a succession of *elementary events*. In our numerical model, elementary events come in one of two basic flavors: *protrusion events* and *retraction events*. During a protrusion event, cell $\alpha$ (referred to as "source cell") incorporates one grid site $\mathbf{x}_t$ ("target grid site") from its neighborhood $\mathcal{N}^{(\alpha)}$,

$$\mathcal{D}^{(\alpha)}_{\text{old}} \to \mathcal{D}^{(\alpha)}_{\text{new}} = \mathcal{D}^{(\alpha)}_{\text{old}} \cup \{\mathbf{x}_t\}, \quad \mathbf{x}_t \in \mathcal{N}^{(\alpha)}, \tag{S5}$$

thereby increasing its cellular domain by one grid site. Here, the neighborhood of cell $\alpha$, $\mathcal{N}^{(\alpha)}$ is defined as

$$\mathcal{N}^{(\alpha)} = \Big\{ \mathbf{x}_l \, \Big| \min_{\mathbf{x}_k \in \mathcal{D}^{(\alpha)}} ||\mathbf{x}_l - \mathbf{x}_k|| = 1 \Big\}. \tag{S6}$$

During a retraction event, source cell $\alpha$ expels one of its membrane grid sites $\mathbf{x}_s \in \mathcal{B}^{(\alpha)}$,

$$\mathcal{D}^{(\alpha)}_{\text{old}} \to \mathcal{D}^{(\alpha)}_{\text{new}} = \mathcal{D}^{(\alpha)}_{\text{old}} \setminus \{\mathbf{x}_s\}, \quad \mathbf{x}_s \in \mathcal{B}^{(\alpha)}_{\text{old}}, \tag{S7}$$

where the set of membrane grid sites $\mathcal{B}^{(\alpha)}$ is defined as

$$\mathcal{B}^{(\alpha)} = \Big\{ \mathbf{x}_k \in \mathcal{D}^{(\alpha)} \, \Big| \min_{\mathbf{x}_l \in \mathcal{N}^{(\alpha)}} ||\mathbf{x}_k - \mathbf{x}_l|| = 1 \Big\}. \tag{S8}$$

Protrusion and retraction events are the numerical analogs of cell protrusions and cell retractions.

In implementing the reassignment rules, Eqs. (S5) and (S7), we have to take into account that cellular domains must not overlap. For solitary cells moving in free space this does not imply any restrictions, and Eqs. (S5) and (S7) apply directly. In the bulk of a confluent monolayer of adhesive cells, however, any protrusion of source cell $\alpha$ into the domain of cell $\beta$ (referred to as "target cell") must be accompanied by a corresponding retraction event $\mathcal{D}^{(\beta)}_{\text{old}} \to \mathcal{D}^{(\beta)}_{\text{new}} = \mathcal{D}^{(\beta)}_{\text{old}} \setminus \{\mathbf{x}_t\}$, where $\mathbf{x}_t$ denotes the target grid site annexed by cell $\alpha$. We emphasize, however, that the reverse is not generally true. If source cell $\alpha$ retracts, i.e. loses one of its boundary grid sites $\mathbf{x}_s \in \mathcal{B}^{(\alpha)}$, the lost grid site $\mathbf{x}_s$ faces either one of two conceivable fates: If, on the one hand, cohesion among cells is sufficiently strong (cf. section S.I D 4 for a definition of the notion "sufficiently strong"), the retraction of cell $\alpha$ exerts a pulling force on one of its neighboring cells $\beta$ (the "target cell"), which is then forced to fill the emerging void at $\mathbf{x}_s$, i.e. $\mathcal{D}^{(\beta)}_{\text{old}} \to \mathcal{D}^{(\beta)}_{\text{new}} = \mathcal{D}^{(\beta)}_{\text{old}} \cup \{\mathbf{x}_s\}$, where $\mathbf{x}_s$ denotes the grid site lost by cell $\alpha$. On the other hand, if adhesion between cells is weak, retraction of the source cell $\alpha$ can lead to a rupture of pre-existing cell contacts between $\alpha$ and other cells at $\mathbf{x}_s$, such that the lost grid site $\mathbf{x}_s$ becomes free space $[c(\mathbf{x}_s) = \alpha \geq 0 \to c(\mathbf{x}_s) < 0]$. Details on the actual implementation of cell rupture are discussed in section S.I D 4.

##### 2. *Monte-Carlo scheme*

In the spirit of a standard Monte-Carlo scheme, the actual simulation proceeds via a succession of *"Monte-Carlo steps"*, where each Monte-Carlo step (MCS) propagates the state of the simulated cell population from time $t$ to time $t+1$. One MCS consists in a series of attempts to perform elementary events, originating from randomly chosen membrane grid sites of randomly chosen cells. The duration of one MCS, i.e. the actual number of attempted elementary events, is chosen such that each of the cells' membrane segments is given the opportunity to attempt, on average, one elementary event per MCS. During each MCS, cell domains $\mathcal{D}^{(\alpha)}$ as well as the numerical values of cell areas $A_\alpha$ and perimeters $P_\alpha$ are updated "on the fly", while the cells' cytoskeletal fields are updated only once at the end of each MCS; cf. section S.I D 2 for the details of this update rule. The simulation then proceeds along the following Monte-Carlo scheme:

1) Initialize the cell population and define the duration of the simulation, i.e. the number of MCS, $N_{\text{mcs}}$, to be performed.

2) Set the simulation time $t = 0$.

3) Perform the next MCS; this step is further detailed below.

4) Update cytoskelatal fields (cf. section S.I D 2).

5) Set $t = t + 1$.

6) Repeat steps 3) – 5) while $t < N_{\text{mcs}}$.

The implementation of a MCS, i.e. the sequence of elementary events is based on the following general considerations:

*(i) Choice of source and target grid sites.* Each elementary event $\mathcal{T}$ originates from a membrane grid site $\mathbf{x}_s \in \mathcal{B}^{(\alpha)}$ of some cell $\alpha$, referred to as *source cell*, which will be referred to as *"source grid site"*. In addition, each elementary event involves a second grid site which lies in the neighborhood of the source grid site $\mathbf{x}_s$ and which is not currently occupied by cell $\alpha$: $\mathbf{x}_t \in \mathcal{N}_s \setminus \mathcal{D}^{(\alpha)}$. In what follows, this additional grid site $\mathbf{x}_t$ will be referred to as *"target grid site"*. This grid site may either be an empty substrate site or a membrane site of another cell $\beta$, in which case the respective cell will be referred to as *target cell*. While the source grid site determines the location of the attempted elementary event, the target grid site determines the direction along which the elementary event is bound to proceed.

*(ii) Monte-Carlo method to generate the system's dynamics.* As mentioned above, the actual dynamics of cells in our computational model is driven by a succession of elementary events, whose cumulative effects over time allow cells to change shapes and to move relative to the substrate as well as relative to each other. Following a standard Monte-Carlo procedure, the probability of occurrence of elementary events $\mathcal{T}$ is determined by a goal function $p(\mathcal{T})$ [cf. point (iii) below]. However, since elementary events come in two basic flavors, protrusions $\mathcal{T}_{\text{pro}}$ and retractions $\mathcal{T}_{\text{ret}}$, their actual occurrence is controlled by a three-step process, once source and target grid sites have been determined: In a first step, the goal function $p$ is used to compute the occurrence probabilities of the two alternative scenarios where either the source cell protrudes toward $\mathbf{x}_t$ [$p(\mathcal{T}_{\text{pro}})$], or retracts from $\mathbf{x}_s$, [$p(\mathcal{T}_{\text{ret}})$]. In a second step, both probabilities $p(\mathcal{T}_{\text{pro}})$ and $p(\mathcal{T}_{\text{ret}})$ are compared and, based on this comparison, a decision is made as to whether one attempts $\mathcal{T}_{\text{pro}}$ or $\mathcal{T}_{\text{ret}}$. The actual decision making process underlying our simulation results is guided by enhancing computational performance and is briefly discussed in point (iv) below. Finally, once a decision has been made on its specific nature, in the third step the elementary event $\mathcal{T}$ is being accepted with probability $p(\mathcal{T})$.

*(iii) Choice of the goal function $p(\mathcal{T})$.* As has been detailed above, we use a goal function $p(\mathcal{T})$ to control the occurrence and acceptance of elementary events $\mathcal{T}$. Following the standard cellular Potts model [1, 2], this goal function takes into account the effects of cell contractility and cell-cell adhesion, using, however, a slightly different implementation; cf. sections S.I D 1 and S.I D 3. In addition, we generalized the definition of the goal function $p(\mathcal{T})$ to explicitly take into account a simplified model of cytoskeletal structures and the ensuing polarization of cells. The actual definition of the goal function will be developed in section S.I D, where, moreover, details concerning the implementation of the cell polarization model will be discussed.

The implementation of a single MCS loop is then given by the following simulation scheme:

3A) Determine the current number of trials per Monte-Carlo step (MCS) $K = \sum_\alpha \sum_{\mathbf{x}_k \in \mathcal{B}^{(\alpha)}} 1$ and set the trial counter $n = 0$.

3B) With equal probability, choose a segment of the cell membrane. Because the cell membrane represents the border between lattice sites occupied by cell $\alpha$ and unoccupied by cell $\alpha$, the chosen membrane segment automatically defines the source grid site $\mathbf{x}_s \in \bigcup_\alpha \mathcal{B}^{(\alpha)}$ and the corresponding target grid site $\mathbf{x}_t \in \mathcal{N}^\alpha \cap \mathcal{N}_s$.

3C) With equal probability, choose whether to attempt a protrusion event ($\mathcal{T}_{\text{pro}}$ or a retraction event ($\mathcal{T}_{\text{ret}}$.

3D) Compute the prospective acceptance probability $p(\mathcal{T}_{\text{pro/ret}})$ corresponding to the attempted event, and decide whether to accept the attempted event on the basis of this probability.

3E) If the attempted elementary event has been accepted, update the cellular domains of source cell $\alpha$ and opponent cell $\beta$; for details see section S.I E.

3F) If $s < K$, set $s \to s + 1$ and then repeat steps 3B) through 3E).

### D. Implementation of cellular traits

In this section, we discuss the various contributions of cellular traits to the overall acceptance probability $p(\mathcal{T})$ of an elementary event $\mathcal{T}$. Specifically, our model takes into account cell contractility, the assembly and disassembly of cytoskeletal structures, cell-cell adhesion, and focal adhesions. We will assume that each of these cellular properties contributes independently to the acceptance probability $p$, such that

$$p = \min\{1,\ p_{\text{cont}} \cdot p_{\text{cyto}} \cdot p_{\text{adh}}\}. \tag{S9}$$

Anticipating our discussions in section S.I D 2, the effects due to focal adhesions have been combined with the effects due to assembly and disassembly of cytoskeletal structures in $p_{\text{cyto}}(\mathcal{T})$. In the following sections, we give detailed discussions for each of these contributions, separately.

#### 1. *Cell contractility*

In biological cells, membrane fluctuations are constrained by elastic forces and contractile cytoskeletal structures, which play a vital role in cell migration [3–5]. In our computational approach, we take cell contractility into account by assigning a contractile "energy"

$$\mathcal{H}_{\text{cont}} = \sum_{\alpha} \left[ \kappa_P^{(\alpha)} \, P_\alpha^2 + \kappa_A^{(\alpha)} \, A_\alpha^2 \right] , \tag{S10}$$

with positive coupling constants $\kappa_A^{(\alpha)}$ and $\kappa_P^{(\alpha)}$ characterizing the contractility of cell $\alpha$; for empty substrate sites ($\alpha < 0$) we set $\kappa_P^{(\alpha)} = \kappa_A^{(\alpha)} = 0$. According to Eq. (S10), the cell's "contractile energy" increases with increasing cell perimeter and increasing cell area. The model Hamiltonian $\mathcal{H}_{\text{cont}}$ can then be used to specify the contractile contribution to the goal function $p(\mathcal{T})$. To this end, let $\Delta\mathcal{H}_{\text{cont}}(\mathcal{T})$ denote the contractile contribution to the energy difference entailed by accepting an elementary event $\mathcal{T}$. Following a standard Metropolis algorithm, we then define

$$p_{\text{cont}}(\mathcal{T}) := \exp\big[-\Delta\mathcal{H}_{\text{cont}}(\mathcal{T})\big]. \tag{S11}$$

The contractile "energy", Eq. (S10), is similar to the corresponding energy model commonly used in cellular Potts models [6]. Unlike the standard cellular Potts model, however, where a target area and target perimeter is used to keep the simulated cells from collapsing, the energetic contribution in Eq. (S10) strictly contracts the cell's body. As will be detailed in the next section, to counteract these contractile forces, we explicitly model cytoskeletal structures within each cell, which provide outward pushing forces to balance cell contraction.

#### 2. *Cytoskeletal structures and focal adhesion*

The cytoskeleton plays key roles both in maintaining the mechanical integrity of the cell and in the process of active cell migration [3, 5, 7]. Our model design aims at achieving high computational efficiency to allow for the simulation of very large cell numbers [8] and, at the same time, to capture the essential effects of cytoskeletal dynamics to attain meaningful results down to the level of single cells. Thus, instead of accounting for a detailed biochemical description by means of reaction-diffusion networks [9, 10], we resort to a simplified implementation of the most pertinent features of cytoskeletal dynamics. Specifically, we propose a rule-based algorithm to model cytoskeletal structures and to assess the integrated effects of cell polarity, cell contractility and adhesion on the collective dynamics of cells as parts of larger groups.

To this end, we define a scalar field $\rho(\mathbf{x}_n)$, $\mathbf{x}_n \in \mathcal{D}^{(\alpha)}$, on the domain of each cell $\alpha$. The local quantity $\rho(\mathbf{x}_n)$ will be referred to as *cytoskelatal field* and is taken to be a measure for the density of cytoskeletal structures at position $\mathbf{x}_n$ within the cell's body. The field variable $\rho(\mathbf{x}_n)$ is dynamically updated as the simulation progresses, reflecting cytoskeletal remodeling. To set up a system of rules underlying the actual implementation of these cytoskeletal remodeling processes, we resort to the following biologically motivated premises:

*(i) The scalar cytoskeletal field $\rho$ is bounded:* The dynamics of cytoskeletal remodeling not only depends on the local number (density) of actin monomers and polymers, but also on a multitude of accessory proteins controlling cytoskeleton assembly and disassembly. Several biological factors, including the action of sequestering proteins like thymosin-$\beta$4, which act to suppress actin polymerization, limited amounts of nucleating proteins like the activated Arp 2/3 complex, and the action of capping proteins keep the local density of actin filaments bounded. We, therefore, introduce bounds for the *cytoskeletal field*: $q \leq \rho(\mathbf{x}_n) \leq Q$. These bounds are cell-type specific. While the upper bound $Q$ mainly reflects the limited availability of protein resources, the lower bound $q$ serves to prevent cells from collapsing.

*(ii) Regulatory proteins affect assembly and disassembly of cytoskeletal structures:* The assembly and disassembly of cytoskeletal structures, numerically encoded by $\rho(\mathbf{x}_n)$, is regulated by a myriad of accessory proteins. In our computational model we simplify these complex processes by resorting to a single "bookkeeping variable" which we will refer to as "regulatory factors". Its local level is stored as an integer variable $F(\mathbf{x}_n)$ for each grid site $\mathbf{x}_n \in \mathcal{D}^{(\alpha)}$. We use $F(\mathbf{x}_n)$ to implement the overall action of regulatory cytoskeletal proteins in an effective and collective manner. Specifically, since the formation of lamellipodial structures depends on active nucleation promoting factors [11], we assume that positive levels, $F(\mathbf{x}_n) > 0$, reflect local conditions in support of network-assembly, whereas non-positive levels, $F(\mathbf{x}_n) \leq 0$, represent predominantly degrading (or disassembly) conditions.

*(iii) Feedback between cytoskeletal structures and regulatory factors:* The activities of accessory cytoskeletal proteins which regulate the local levels of cytoskeletal structures are themselves controlled by a number of mechanical and chemical signals received by the cell. Here and in the following, our focus will be on mechanical signals. For example, important regulatory proteins like the Arp 2/3 complex are activated locally at the cell membrane, from where they diffuse into the bulk of the cell until they are bound by actin [11–13]. Adopting a coarse level of description, this diffusion-degradation dynamics entails a finite range of regulatory proteins, which are activated at the cell's membrane. In our model, we use the integer variable variable $F(\mathbf{x}_n) > 0$ to implement this propagation of mechanical information, perceived by cell $\alpha$ at its periphery $\mathcal{B}^{(\alpha)}$, across a certain spatial distance $R$. The local levels of $F(\mathbf{x}_n)$ are continuously updated as the MCS loop progresses. The actual update procedure is given by the following set of rules; cf. Fig. S2:

- If a *protrusion event* has been accepted at the source site $\mathbf{x}_s \in \mathcal{B}^{(\alpha)}$ (source cell: $\alpha$; target cell: $\beta$), then for all sites $\mathbf{x}_n$ within a range $R$ (i.e. $||\mathbf{x}_n - \mathbf{x}_s|| < R$) the integer variable signifying regulatory factors is incremented up and down for the

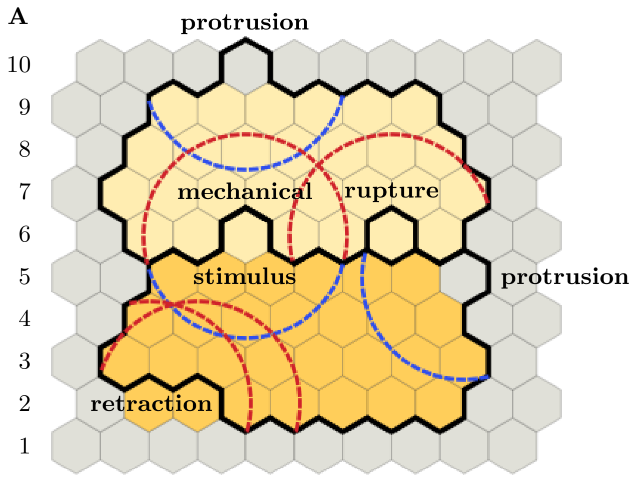


FIG. S2. Distribution of regulatory factors on the basis of accepted elementary events. For ease of reference, grid rows have been numbered from 1 to 10. *Left*: Solid black line indicate cells' membrane positions *after* acceptance of the respective elementary event; colors indicate cellular domains *before* the respective elementary event has been accepted (gray: substrate; shades of yellow: cells). Blue and red circular arcs (of radius $R$) delineate areas of local increase or decrease in the level of regulatory factors, respectively. The following elementary events are depicted: (i) lower cell retracts (two grid sites in row 2); (ii) lower cell protrudes (row 5); (iii) upper cell protrudes (row 10). In addition, the following elementary events occur across the cell-cell boundary: (iv) retraction of upper cell leads to rupture of cell-cell contact (row 6, right event); (v) either the lower cell protrudes and pushes the upper cell or the upper cell retracts and pulls on the lower cell (row 6, left event). Specifically, event (v) entails mechanical signaling between the upper and lower cell and, therefore, affects the distribution of regulatory factors in *both* cells. *Right*: Identical copy of the left image. Colors indicate local levels of regulatory factors $F$ (blue: $F$ positive; white: $F$ zero; red: $F$ negative; gray: substrate site). Note, in particular, that a substrate grid site has been inserted where cell rupture occurred (row 6, right pixel). The following cases can be distinguished: (i) Grid site $\mathbf{x}_k$ lies in the zone of influence of only positive (blue circles) or negative (red circles) chemical feedback, in which case the level of regulatory factors is positive or negative, respectively (e.g. red grid sites in row 2, or blue grid sites in row 5). (ii) Grid site $\mathbf{x}_k$ lies outside of any zone of influence, in which case the level regulatory factors is zero (e.g. white grid sites in row 2). (iii) Grid site $\mathbf{x}_k$ lies in the zone of influence of equally many positive and negative feedbacks, in which case the level of regulatory factors remains zero (e.g. fourth grid sites in row 4). (iv) Grid site $\mathbf{x}_k$ lies in a zone of predominantly positive or negative feedback, in which case the level of regulatory factors is positive or negative, respectively (e.g. third grid site in row 4). Recall that only the sign of $F$ is of significance to update the cells' cytoskelatal fields; cf. Eq. (S13).

protruding and the retracting cell, respectively:

$$F(\mathbf{x}_n) \to \begin{cases} F(\mathbf{x}_n) + 1, & \mathbf{x}_n \in \mathcal{D}^{(\alpha)}, \\ F(\mathbf{x}_n) - 1, & \mathbf{x}_n \in \mathcal{D}^{(\beta)}. \end{cases} \quad \text{(S12a)}$$

- Similarly, if a *retraction event* has been accepted at the source site $\mathbf{x}_s \in \mathcal{B}^{(\alpha)}$, and the (local) cell contact between source cell $\alpha$ and target cell $\beta$ has remained intact, then within a range $R$ one applies the inverse update rule:

$$F(\mathbf{x}_n) \to \begin{cases} F(\mathbf{x}_n) - 1, & \mathbf{x}_n \in \mathcal{D}^{(\alpha)}, \\ F(\mathbf{x}_n) + 1, & \mathbf{x}_n \in \mathcal{D}^{(\beta)}. \end{cases} \quad \text{(S12b)}$$

- If a *retraction event* has been accepted at the source site $\mathbf{x}_s \in \mathcal{B}^{(\alpha)}$, and in addition the (local) cell contact between source cell $\alpha$ and target cell $\beta$ has *ruptured*, then the regulatory factors are reduced only within a range R in the retracting cell:

$$F(\mathbf{x}_n) \to \begin{cases} F(\mathbf{x}_n) - 1, & \mathbf{x}_n \in \mathcal{D}^{(\alpha)}, \\ \text{no update}, & \text{else}. \end{cases} \quad \text{(S12c)}$$

Finally, if the target grid site $\mathbf{x}_t$ is not occupied by any cell ($\beta \geq 0$) prior to the elementary event, only the first lines in the above update scheme apply.

By virtue of the above update scheme, Eq. (S12), "regulatory factors" are continuously distributed across each cell's domain $\mathcal{D}^{(\alpha)}$ as the current MCS progresses. At the end of each MCS, the accumulated (local) values of $F(\mathbf{x}_n)$ are used to update the local values of the cytoskelatal fields $\rho(\mathbf{x}_n)$ inside each cell $\alpha \geq 0$ ($\mathbf{x}_n \in \mathcal{D}^{(\alpha)}$): We assume that for positive values, $F > 0$, there is assembly of cytoskeletal structures and $\rho$ is increased by an amount proportional to the distance of $\rho$ from its upper bound $Q$:

$$\rho(\mathbf{x}_n, t + \Delta t) = \rho(\mathbf{x}_n, t) + \mu\,[Q - \rho(\mathbf{x}_n, t)]. \quad \text{(S13a)}$$

Thereby $Q$ is a fixed point of this map and limits the build-up of cytoskeletal structures. In contrast, for negative values, $F(\mathbf{x}_n) \leq 0$, disassembly prevails, and we assume that $\rho$ then tends towards its lower bound $q$:

$$\rho(\mathbf{x}_n, t + \Delta t) = \rho(\mathbf{x}_n, t) + \mu\,[q - \rho(\mathbf{x}_n, t)]. \quad \text{(S13b)}$$

The parameter $\mu$ signifies the rate at which cytoskeletal structures respond to the regulatory factors $F$. For the parameters and cell sizes used in this work ($q, Q = \mathcal{O}(100)$, and each cell occupying $\sim$1000 grid sites) we set $\mu = 0.1$.

After this update procedure for $\rho(\mathbf{x}_n, t)$ is completed, all regulatory factors are reset, $F(\mathbf{x}_n) = 0, \forall n$. This prevents "spurious memory effects" which may arise once the cell's rear reaches its initial leading edge position as time goes on. In essence, resetting regulatory factors upon completion of one MCS implies that the diffusion-degradation dynamics, underlying the distribution of regulatory factors, is fast on the scale of one MCS.

We emphasize that the cytoskelatal field $\rho(\mathbf{x}_n)$ is defined only for grid sites $\mathbf{x}_n \in \mathcal{D}^{(\alpha)}$ occupied by an actual cell ($\alpha \geq 0$). To allow for spatial variations of substrate properties, we therefore introduce a second scalar field variable $\varphi(\mathbf{x}_n)$, which is defined on the entire computational grid. The scalar field $\varphi(\mathbf{x}_n)$ is taken to measure the local density of anchoring points that a cell might use to form focal adhesions. Although one might consider to treat $\varphi$ as a time-dependent field variable, in this work $\varphi$ is used to implement static substrate patterns, only. The field $\varphi(\mathbf{x}_n)$ is thus initialized once at the beginning of the simulation and kept fixed throughout the entire simulation.

Having introduced the fields $\rho(\mathbf{x}_n)$ and $\varphi(\mathbf{x}_n)$, we now discuss their impact on the system's dynamics by giving their contribution to the goal function $p(\mathcal{T})$. Assume that the elementary event $\mathcal{T}$ is attempted by a source cell $\alpha$ at source grid site $\mathbf{x}_s \in \mathcal{B}^{(\alpha)}$. Further, let $\mathbf{x}_t$ denote the target grid site and $\beta$ denote the index of the target cell (where as ususal $\beta \geq 0$ indicates that $\mathbf{x}_t$ is occupied by an actual cell, $\beta < 0$ means that $\mathbf{x}_t$ exposes substrate). We then define the *"polarization energy"* $\Delta\mathcal{H}_{\text{cyto}}(\mathcal{T})$ as follows:

$$\Delta\mathcal{H}_{\text{cyto}}(\mathcal{T}) \equiv \begin{cases} \rho(\mathbf{x}_t) - \rho(\mathbf{x}_s), & \mathcal{T} \,\hat{=}\, \mathcal{T}_{\text{pro}} \wedge \beta \geq 0, \\ \rho(\mathbf{x}_s) - \rho(\mathbf{x}_t), & \mathcal{T} \,\hat{=}\, \mathcal{T}_{\text{ret}} \wedge \beta \geq 0, \\ -\big[\rho(\mathbf{x}_s) + \varphi(\mathbf{x}_t)\big], & \mathcal{T} \,\hat{=}\, \mathcal{T}_{\text{pro}} \wedge \beta < 0, \\ \rho(\mathbf{x}_s) + \varphi(\mathbf{x}_t), & \mathcal{T} \,\hat{=}\, \mathcal{T}_{\text{ret}} \wedge \beta < 0, \end{cases} \tag{S14a}$$

Here, the definition of $\Delta\mathcal{H}_{\text{cyto}}$ is such that the likelihood of cell protrusions is enhanced if the concentration of cytoskeletal structures at the source grid site, $\rho(\mathbf{x}_s)$, is larger than the concentration at the target grid site, $\rho(\mathbf{x}_t)$ [first row of Eq. (S14a)], and vice versa for cell retractions [second row of Eq. (S14a)]. The strength of focal adhesions is taken to be measured by the sum $\rho + \varphi$. Their associated "anchoring effects" (which increase with growing strength of focal adhesions) promote the formation of cell protrusions against unoccupied substrate sites [third row of Eq. (S14a)], and, correspondingly, impedes cell retractions [fourth row of Eq. (S14a)]. Note, in particular, that the first two rows of Eq. (S14a) can be obtained from a combined evaluation of the lower two rows. For example, if source cell $\alpha$ annexes $\mathbf{x}_s$ starting from $\mathbf{x}_t$, two things need to happen: First, focal adhesions formed by the target cell $\beta$ must be broken, implying a contribution $\Delta\mathcal{H}_{\text{cyto}} = \rho(\mathbf{x}_t) + \varphi(\mathbf{x}_t)$ [fourth row of Eq. (S14a)]. Secondly, new focal adhesions are formed by the source cell $\alpha$, implying a contribution $\Delta\mathcal{H}_{\text{cyto}} = -[\rho(\mathbf{x}_s) + \varphi(\mathbf{x}_t)]$ [third row of Eq. (S14a)]. Taking the sum of both contributions gives the expression in the first row of Eq. (S14a). An analogous line of arguments leads to the expression in the second row of Eq. (S14a).

The contribution to the goal function $p(\mathcal{T})$ due to the *polarization energy* $\Delta\mathcal{H}_{\text{cyto}}(\mathcal{T})$ is then defined by

$$p_{\text{cyto}}(\mathcal{T}) \equiv \exp\big[-\Delta\mathcal{H}_{\text{cyto}}(\mathcal{T})\big], \tag{S14b}$$

where the characteristic "energy scale" for $\Delta\mathcal{H}_{\text{cyto}}$ is set by the polarization bounds $q$ and $Q$, which turns out to have important implications for collective cell dynamics, as discussed in the main text.

#### 3. *Cell adhesion*

To implement the ability of cells to establish cell adhesions across cell-cell interfaces, we use a special form for the respective contribution to the goal function $p$, which is designed to distinguish between the formation of new and the breakage of existing cell-cell adhesion sites.

To this end, we define *adhesion matrices* $B_{\alpha,\beta}$ and $B'_{\alpha,\beta}$ quantifying the system's change in "energy" upon forming a new contact between cells $\alpha$ and $\beta$ [$B_{\alpha,\beta}$] and upon breaking a pre-existing contact between those cells by an "intruder cell" $\gamma \neq \alpha, \beta$ [$B'_{\alpha,\beta}$]. In our computational model, we assume that formation of new cell-cell contacts is energetically favored, and that breaking of pre-existing contacts by intruder cells is energetically penalized. The matrix entires of $B_{\alpha,\beta}$ and $B'_{\alpha,\beta}$, therefore, have a definite sign, which we take to be positive. The ordering of the cell index pair of $B_{\alpha,\beta}$ and $B'_{\alpha,\beta}$ is of no physical significance, i.e. the adhesion matrices are symmetric. We also assume that a given cell $\alpha$ does not interact with itself, such that the diagonal elements of the adhesion matrices vanish. Finally, there is no adhesion between cells and empty substrate sites, such that all matrix elements containing a negative cell index vanish. In summary, the adhesion matrices $B_{\alpha,\beta}$ and $B'_{\alpha,\beta}$ exhibit the following properties:

$$B_{\alpha,\beta} = B_{\beta,\alpha} \geq 0, \tag{S15a}$$
$$B'_{\alpha,\beta} = B'_{\beta,\alpha} \geq 0, \tag{S15b}$$
$$B_{\alpha,\alpha} = B'_{\alpha,\alpha} = 0, \tag{S15c}$$
$$B_{\alpha,\beta} = B'_{\alpha,\beta} = 0, \qquad \text{if } \alpha < 0 \vee \beta < 0. \tag{S15d}$$

In addition, we assume that the energy cost associated with the breakage of a given cell-cell contact exceeds the energetic benefit of its initial formation, i.e.

$$B'_{\alpha,\beta} \geq B_{\alpha,\beta}, \tag{S15e}$$

where equality of both quantities would reproduce the assumption underlying the standard cellular Potts model [1, 2]. We shall refer to this property as the "dissipative nature of cell-cell adhesion".

To implement the effects of cell-cell adhesion, we compute the "energy difference" $\Delta\mathcal{H}_{\text{adh}}(\mathcal{T})$ for any given elementary event $\mathcal{T}$ according to the scheme illustrated in Fig. S3. One has to distinguish between *protrusion* and *retraction* events. First, say that a cell $\alpha$ attempts a protrusion event $\mathcal{T}_{\text{pro}}$, involving the source grid site $\mathbf{x}_s \in \mathcal{B}^{(\alpha)}$ and the target grid site $\mathbf{x}_t \in \mathcal{B}^{(\beta)}$, as illustrated in Fig. S3A. In this case, cell $\alpha$ acts as intruder cell, since the depicted protrusion event affects three pre-existing contacts between the target cells $\beta$ and a "third party" cell $\gamma$. Acceptance of the depicted protrusion event would have the following energetically relevant effects: (i) All pre-existing contacts between the target cell $\beta$ and third party cell $\gamma \neq \alpha, \beta$ at the target grid site $\mathbf{x}_t$ are torn apart. (ii) New contacts between the source cell $\alpha$ and third party cell $\gamma \neq \alpha, \beta$ are being established. (iii) The length of the cell contact line between source cell $\alpha$ and target cell $\beta$ is changed. Altogether, these three effects lead to the following cell adhesion energy difference,

$$\Delta\mathcal{H}_{\text{adh}}(\mathcal{T}_{\text{pro}}) \equiv -\sum_{\mathbf{x}_j \in \mathcal{N}_t} \left[B_{\alpha,c(\mathbf{x}_j)} - \delta_{\alpha,c(\mathbf{x}_j)}\, B_{\beta,c(\mathbf{x}_j)}\right] + \sum_{\mathbf{x}_j \in \mathcal{N}_t} B'_{\beta,c(\mathbf{x}_j)}(1-\delta_{\alpha,c(\mathbf{x}_j)}). \tag{S16a}$$

The first term in this expression accounts for the (energetically favored) formation of new cellular contacts, as well as for the remodeling of the interface between source cell $\alpha$ and target cell $\beta$ [points (ii) and (iii)]. The second term measures the (energetically penalized) breaking of pre-existing cell contacts [point (i)] and, therefore, impedes cell $\alpha$'s ability to intrude. Conversely, if source cell $\alpha$ attempts a retraction event $\mathcal{T}_{\text{ret}}$, the same reasoning leading to Eq. (S16a) applies, only this time the elementary event proceeds in reverse, i.e. from the target site $\mathbf{x}_t$ to the source site $\mathbf{x}_s$; cf. Fig. S3A:

$$\Delta\mathcal{H}_{\text{adh}}(\mathcal{T}_{\text{ret}}) \equiv -\sum_{\mathbf{x}_j \in \mathcal{N}_s} \left[B_{\beta,c(\mathbf{x}_j)} - \delta_{\beta,c(\mathbf{x}_j)}\, B_{\alpha,c(\mathbf{x}_j)}\right] + \sum_{\mathbf{x}_j \in \mathcal{N}_s} B'_{\alpha,c(\mathbf{x}_j)}(1-\delta_{\beta,c(\mathbf{x}_j)}). \tag{S16b}$$

We may now use Eqs. (S16)(a,b) to illustrate the "dissipative nature" of adhesive interactions by means of two archetypical examples. First, consider the adhesive energy contribution to any cyclic process. By a cyclic process we mean a sequence of two mutually inverse elementary events, e.g. a protrusion event $\mathcal{T}_{\text{pro}}$, which is immediately followed by its inverse retraction event $\mathcal{T}_{\text{ret}}$, such that the system's final configuration is identical to its initial configuration. Using Eqs. (S16) we find for the total adhesive energy contribution to a cyclic process:

$$\Delta\mathcal{H}^{(\text{cycl})}_{\text{adh}} = \sum_{\mathbf{x}_j \in \mathcal{N}_t \setminus \left(\mathcal{D}^{(\alpha)} \cup \mathcal{D}^{(\beta)}\right)} \left[(\Delta B)_{\alpha,c(\mathbf{x}_j)} + (\Delta B)_{\beta,c(\mathbf{x}_j)}\right], \tag{S17}$$

$$(\Delta B)_{\alpha,\beta} := B'_{\alpha,\beta} - B_{\alpha,\beta} \geq 0 \quad [\text{Eq. (S15e)}], \tag{S18}$$

and can therefore conclude that

$$\Delta\mathcal{H}^{(\text{cycl})}_{\text{adh}} \geq 0,$$

where $\mathcal{N}_t$ denotes the neighborhood of the grid site which temporarily changes its cell index, and where $\alpha$ and $\beta$ are the indices of the source and target cells involved in the cyclic process; cf. Fig. S3A. Since $\Delta_{\alpha,\beta} \geq 0$, the above adhesive energy contribution is non-negative, thus leading to an amount of energy equal to $\Delta\mathcal{H}^{(\text{cycl})}_{\text{adh}}$ being dissipated as the cyclic process completes. This leads us to refer to the parameter matrix $\Delta_{\alpha,\beta}$ as *"dissipation matrix"*. Second, consider two (infinitely extended) rows of cells in adhesive contact, sliding past each other. This situation is depicted in Fig. S3B, where the top row of cells moves (as a whole) to the right by one grid site, while the bottom row of cells remains stationary. To assess the adhesive energy contribution along the contact line connecting both cell rows, note that the depicted transformation can be implemented by letting each cell in the top row protrude its leading (i.e. right) edge by one grid site. For each protruding (source) cell $\alpha$, this transformation entails to the following energetic effects (cf. discussion above): (i) Two of the pre-existing cell-cell contacts between the source cell's right neighbor in the top row (target cell $\beta$) and the corresponding cell in the bottom row (third party cell $\gamma$) get torn apart, leading to an energetic contribution $2\,B_{\beta,\gamma}$. (ii) In return, two new contacts between the protruding (source) cell $\alpha$ and cell $\gamma$ are being established, leading to a contribution $-2\,A_{\alpha,\gamma}$. (iii) Since the length of the contact line between cells in the top row (i.e. between protruding source cell $\alpha$ and retracting target cell $\beta$) remains unchanged, there's no further energetic contribution due to adhesive contacts between cells in the top row. Assuming that all cells in the system are of equal types, we write $A_{\alpha,\beta} \equiv A$ and $B_{\alpha,\beta} \equiv B$ $(\alpha \neq \beta)$ and, therefore, find

$$\Delta\mathcal{H}^{(\text{visc})}_{\text{adh}} = 2(B-A) \equiv 2\Delta \geq 0, \tag{S19}$$

i.e. a non-negative dissipative contribution per cell. The size of the dissipation parameter $\Delta$ thus introduces a natural means to tune the system's *shear viscosity*.

With the above definitions of the adhesive energy changes, Eqs. (S16), we define the contribution of cell adhesion to the goal function $p(\mathcal{T})$ as follows:

$$p_{\text{adh}}(\mathcal{T}) \equiv \exp\left[-\Delta\mathcal{H}_{\text{adh}}(\mathcal{T})\right]. \tag{S20}$$

#### 4. *Rupture of cell contacts*

By now, we have introduced all components making up the total acceptance probability $p(\mathcal{T})$, Eq. (S9). To conclude our discussions concerning the implementation of cellular traits, we highlight one additional aspect of elementary events. So far, the notion of an elementary event can be summarized as follows: Once source and target grid sites, $\mathbf{x}_s$ and $\mathbf{x}_t$, have been selected, acceptance of a protrusion [retraction] event causes (among

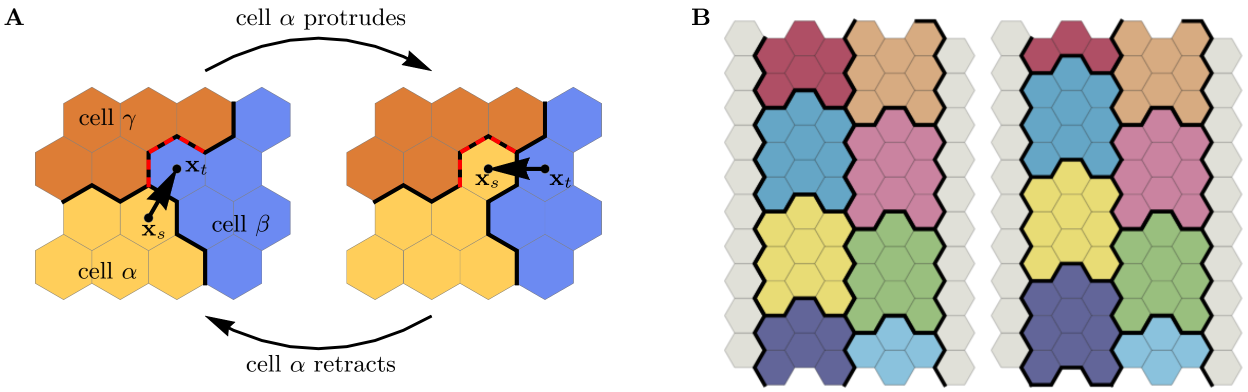


FIG. S3. Cell-cell adhesion. **A)** Adhesive energy contribution in a cyclic process, where a protrusion of source cell $\alpha$ against target cell $\beta$ is followed by the inverse retraction event. Both events involve a third party cell $\gamma$, leading to net energy dissipation after the cyclic process has been completed. *Protrusion:* (i) Three pre-existing cell-cell contacts between $\beta$ and $\gamma$ are torn apart (red dashed contacts); (ii) three new contacts between $\alpha$ and $\gamma$ are formed; (iii) the contact length between source cell $\alpha$ and target cell $\beta$ increases by one unit of length. This implies $\Delta\mathcal{H}_{\text{adh}}(\mathcal{T}_{\text{pro}}) = 3B_{\beta,\gamma} - 3A_{\alpha,\gamma} - A_{\alpha,\beta}$. *Retraction:* (i) Three pre-existing cell-cell contacts between $\alpha$ and $\gamma$ are torn apart (red dashed contacts); (ii) three new contacts between $\beta$ and $\gamma$ are formed; (iii) the contact length between source cell $\alpha$ and target cell $\beta$ decreases by one unit of length. This implies $\Delta\mathcal{H}_{\text{adh}}(\mathcal{T}_{\text{ret}}) = 3B_{\alpha,\gamma} - 3A_{\beta,\gamma} + A_{\alpha,\beta}$. Altogether, this leads to $\Delta\mathcal{H}^{(\text{cycl})}_{\text{adh}} = \Delta\mathcal{H}_{\text{adh}}(\mathcal{T}_{\text{pro}}) + \Delta\mathcal{H}_{\text{adh}}(\mathcal{T}_{\text{ret}}) = 3\Delta_{\alpha,\gamma} + 3\Delta_{\beta,\gamma} \geq 0$, i.e. a (non-negative) dissipative contribution, whose magnitude depends on the dissipation matrix $\Delta_{\alpha,\beta} = B_{\alpha,\beta} - A_{\alpha,\beta} \geq 0$. **B)** Shear viscosity due to cell-cell adhesion. Consider two rows of adhesive cells sliding past each other as indicated in the figure (left row of cells moves up by one gird site; colors indicate different cells). The associated adhesion energy (per cell) reads $\Delta\mathcal{H}_{\text{adh}}/n_c = 2\,(B - A) \geq 0$, where $n_c$ denotes the number of cells sliding past each other, and where we assumed cells of like type, i.e. $A_{\alpha,\beta} \equiv A$ and $B_{\alpha,\beta} \equiv B$ $(\alpha \neq \beta)$. The condition $B > A$, Eq. (S15e), thus implies positive friction associated with cellular shear flows, whose magnitude is proportional to the number of cells sliding past each other. Note that this shear viscosity vanishes for $A = B$, i.e. for zero dissipation matrix.

other things like the distribution of regulatory factors) the cell index to be copied from $\mathbf{x}_s$ to $\mathbf{x}_t$ [from $\mathbf{x}_t$ to $\mathbf{x}_s$]. In other words, the domain $\mathcal{D}^{(\alpha)}$ of source cell $\alpha$ annexes $\mathbf{x}_t$ [loses $\mathbf{x}_s$], while the domain $\mathcal{D}^{(\beta)}$ of source cell $\beta$ is forced to let go $\mathbf{x}_t$ [accommodate $\mathbf{x}_s$]. However, if both, source and target cells are actual cells, i.e. $\alpha, \beta \geq 0$, and if the source cell attempts a retraction event, there is one additional possible outcome: If cell cohesion is weak, the pulling force exerted by the retracting source cell $\alpha$ on its neighboring cells might also result in rupture of all pre-existing contacts between the retracting source cell and its neighboring cells at $\mathbf{x}_s$, rather than forcing one of its neighboring cells (the target cell) to fill the void created at $\mathbf{x}_s$ once $\alpha$ retracts; cf. rupture event depicted in Fig. S2. To test for the occurrence of cell rupture, the total energetic cost of each attempted retraction event between two actual cells is evaluated under two different assumptions: First, we assume that the pulling force exerted by the retracting source cell $\alpha$ on target cell $\beta$ is strong enough to force $\beta$ to fill the void created at $\mathbf{x}_s$ (i.e. to accommodate $\mathbf{x}_s$), and call this a regular retraction event $\mathcal{T}_{\text{ret}}$. Secondly, we assume that the retraction of source cell $\alpha$ causes all pre-existing cell-cell contacts of cell $\alpha$ at $\mathbf{x}_s$ to rupture, leaving a free substrate site at $\mathbf{x}_s$ after the retraction event has been accepted. This latter event will be referred to as rupture event $\mathcal{T}_{\text{rup}}$. We then compute the total energy differences

$$\Delta\mathcal{H}(\mathcal{T}_{\text{ret}}) = \Delta\mathcal{H}_{\text{el}}(\mathcal{T}_{\text{ret}}) + \Delta\mathcal{H}_{\text{cyto}}(\mathcal{T}_{\text{ret}}) + \Delta\mathcal{H}_{\text{adh}}(\mathcal{T}_{\text{ret}})$$

and

$$\Delta\mathcal{H}(\mathcal{T}_{\text{rup}}) = \Delta\mathcal{H}_{\text{el}}(\mathcal{T}_{\text{rup}}) + \Delta\mathcal{H}_{\text{cyto}}(\mathcal{T}_{\text{rup}}) + \Delta\mathcal{H}_{\text{adh}}(\mathcal{T}_{\text{rup}})$$

under both assumptions [14] and compare the respective outcomes. Then, if the rupture event is energetically favored over the regular retraction event, i.e. $\Delta\mathcal{H}(\mathcal{T}_{\text{rup}}) < \Delta\mathcal{H}(\mathcal{T}_{\text{ret}})$, cohesion between cells is weak. In this case, the rupture event $\mathcal{T}_{\text{rup}}$, rather than the regular retraction event $\mathcal{T}_{\text{ret}}$ will be attempted. Otherwise, cohesion is strong and a regular retraction event $\mathcal{T}_{\text{ret}}$ will be attempted.

#### 5. *Dissipation in substrate adhesion*

We introduce dissipation in substrate adhesion by leaving the Hamiltonian unaltered for protrusion events but adding a term for retraction events:

$$\Delta\mathcal{H}(\mathcal{T}_{\text{ret}}) \to \Delta\mathcal{H}(\mathcal{T}_{\text{ret}}) + D. \qquad \text{(S21)}$$

A cell that adheres to the substrate at some grid point has to pay a cost to retract from it. In other words, we assume that an energy $D$ is dissipated once the adhesive bonds between the cell and the substrate break. We study $D \in [0, \Delta Q]$.

To keep its overall size across translations, the cell has to gain and lose equal amounts of hexagons, with $\Delta Q = Q - q$ as the maximal energy gained by a single gain-and-loss in the absence of dissipation. In the presence of dissipation however, the cell has to pay at least a cost of $q + D$ to detach at an arbitrary location, resulting in $(Q - q) - D$ as the maximal energy gained by a single gain-and-loss in the presence of dissipation. Thus, while for $D = 0$ there is no impact of substrate dissipation on cell motility, it will at the latest for $D = \Delta Q$ completely inhibit (directed) cell migration.

#### E. Cell domain update routine

Having discussed the implementation concerning the two basic types of elementary events, viz. protrusion events $\mathcal{T}_{\text{pro}}$ and retraction events $\mathcal{T}_{\text{ret}}$, as well as the two subtypes of regular retraction events and rupture events $\mathcal{T}_{\text{rup}}$, we can now summarize the cell domain update routine, point 3.3E) in section S.I C 2. To this end, and in accordance with our previous notation, we use the cell indices $\alpha$ and $\beta$ to denote source and target cell, and $\mathbf{x}_s$ and $\mathbf{x}_t$ to denote source and target grid site. Moreover, equal signs "=" in the following listing are to be interpreted as assignment operators, where the value of the variable on the left hand side of the operator is assigned to the value of the variable on the right hand side. With these preliminary remarks in mind, the cell update routine can be summarized as follows:

- If the accepted elementary event is a protrusion event:
  1. Set $\rho(\mathbf{x}_t) = \rho(\mathbf{x}_s)$ and $F(\mathbf{x}_t) = F(\mathbf{x}_s)$.
  2. Set $\mathcal{D}^{(\alpha)} \to \mathcal{D}^{(\alpha)} \cup \{\mathbf{x}_t\}$.
  3. Set $\mathcal{D}^{(\beta)} \to \mathcal{D}^{(\beta)} \setminus \{\mathbf{x}_t\}$.
  4. Distribute regulatory factors according to Eq. (S12a).
- If the accepted elementary event is a regular retraction event:
  1. Set $\rho(\mathbf{x}_s) = \rho(\mathbf{x}_t)$ and $F(\mathbf{x}_s) = F(\mathbf{x}_t)$.
  2. Set $\mathcal{D}^{(\alpha)} \to \mathcal{D}^{(\alpha)} \setminus \{\mathbf{x}_s\}$
  3. Set $\mathcal{D}^{(\beta)} \to \mathcal{D}^{(\beta)} \cup \{\mathbf{x}_s\}$
  4. Distribute regulatory factors according to Eq. (S12b).
- If the accepted elementary event is a rupture event:
  1. Set $\rho(\mathbf{x}_s) = 0$ and $F(\mathbf{x}_s) = 0$.
  2. Set $\mathcal{D}^{(\alpha)} \to \mathcal{D}^{(\alpha)} \setminus \{\mathbf{x}_s\}$
  3. Distribute regulatory factors according to Eq. (S12c).

#### F. Cell proliferation and mitosis

While cell proliferation and mitosis play no role in the experimental setup of rotating cell clusters, cell growth and division are observed experimentally in a setup where a sheet of cells expands into free space after removal of a stencil. Therefore, it is essential to include proliferation of cells in the numerical model. How this is done is described in this section.

We distinguish between two phases in the cell cycle, an *interphase* during which cells roughly double in volume and *mitosis*, the process of cell division. Even though a further partitioning of the interphase was considered in previous work [15], we do not expect that such a distinction is relevant for our results. In our computational framework cell growth may be implemented by progressively changing any cellular parameter that affects the cell's equilibrium size. The two possible, largely equivalent choices are a successive reduction of the area coupling constant $\kappa_A^{(\alpha)}$ or an increase of the average cell polarization $(Q^{(\alpha)} + q^{(\alpha)})/2$. We here employ the first method. We assume that individual cells grow exponentially [16] over a well-defined period $T_g$. Additional variability in cell cycle length can be achieved by introducing an additional refractory phase with exponentially distributed waiting times and the average waiting time $T_r$, which we set to $T_r = 0$ in this work. Moreover, we assume that the migratory behavior of a cell should not change significantly as it grows. However, as the cell grows in size by a factor of 2, it also increases its perimeter and the corresponding energy cost for adding new membrane segments roughly by a factor of $\sqrt{2}$. Therefore, as we do not scale the cytoskeletal density $\rho$ and the resulting energy gains for protrusions during cell growth, we mitigate the increased cost for ruffling the membrane by reducing the perimeter stiffness by a factor of $\sqrt{2}$. The quantitative viability of this approach is further discussed in section S.II A.

To prevent tissue overgrowth, cell proliferation is generally contact inhibited in healthy cells: When the tissue approaches a state where each cell has formed adhesive contacts with the substrate and is completely surrounded by neighbours, cells stop proliferating. In addition it has been proposed that the pressure or local density in the tissue has a negative impact on the local growth rate [17, 18]. To account for these phenomena in the model, we complement the two cell cycle periods interphase and mitosis by a quiescent cell state during which cell growth is halted. The parameters $\kappa_A$ and $\kappa_P$ are, therefore, kept constant for a quiescent cell; we denote the corresponding values $\kappa_{A,0}$ and $\kappa_{P,0}$. There are many possible ways to implement contact inhibition in our computational model. For example, it could be implemented by allowing a quiescent cell to enter the cell cycle triggered by low local cell density, or when a sufficiently large fraction of its membrane length is not in contact with neighbour cells but exposed to free space. In our model it proves numerically advantageous to make a quiescent cell enter the interphase when its area succeeds a certain reference area. We choose this area threshold as a fraction $r = 1$ of the equilibrium cell size $A_{\mathrm{ref}}$ reached by a free, solitary cell with constant cytoskeletal density $(Q + q)/2$. Cells living in a densely packed environment will not exceed the area threshold due to the pressure exerted on them by neighboring cells and can, therefore, not grow. Conversely, cells exposed to free space are more likely to reach this threshold and proliferate. Finally, a growing cell in interphase becomes mitotic after the growth time $T_g$ has passed, at which point cell size has roughly doubled with respect to the size in the quiescent period. We assume that cell migration and mitosis are processes that exclude each other. Hence, the positive feedback leading to persistent cell migration is switched off for mitotic cells and the cytoskeletal density relaxes to the neutral state $(Q + q)/2$ according to Eq. 3c.

There appears to be no universal set of rules which determine the orientation of the cleavage plane along which cells divide [19]. Rather, for epithelial tissue there are a variety of factors which include local cell geometry and the direction of stress in tissue [20]. Though it is in principle possible to implement any given rule in our computational model, in its present version the axis along which the cell divides in chosen as a random direction through the geometric center of the cell. In case of irregular cell shapes a separation of the cellular domain into more than two connected components can occur. To prevent violation of topological constraints, in this case the two largest components are considered as descendant cells and the residual pixels are replaced by substrate pixels.

We explicitly account for the finite duration of the mitotic phase $T_m$ by keeping the cells in a mitotic state for the aforementioned time period, until the final instantaneous splitting of the cellular domains. After cell division, persistent cell migration of the daughter cells is enabled again. The descendent cells will subsequently re-enter the growing phase if their area exceeds the defined threshold, as mentioned above.

The following list summarizes the steps motivated and explained in the previous paragraphs. These additional steps are performed in a simulation that includes cell proliferation:

- Assign a state variable $s^{(\alpha)}$ to each cell which encodes the current phase in the cell cycle:

$$s^{(\alpha)} = \begin{cases} 0, \text{ quiescent phase} \\ 1, \text{ refractory phase} \\ 2, \text{ interphase} \\ 3, \text{ mitotic phase} \end{cases} \tag{S22}$$

- Compute the equilibrium size $A_{\mathrm{ref}} = (q + Q - 4\pi\sqrt{3}\kappa_P)/(2\sqrt{3}\kappa_A)$ of a free, solitary cell with fixed cytoskeletal density $\rho = (Q + q)/2$ on the substrate used in the simulation.

- At simulation start $t = 0$, all cells are in the quiescent state, $s^{(\alpha)}(0) = 0$, and start with $Q^{(\alpha)}(0) = Q_0$.

- After completion of each Monte Carlo time step $t$, perform one of the following changes for each cell:

  – Switch from quiescent to refractory state:

$$\begin{aligned} &s^{(\alpha)}(t) = 0 \wedge A^{(\alpha)}(t) > rA_{\mathrm{ref}} \\ &\Rightarrow s^{(\alpha)}(t + 1) = 1. \end{aligned} \tag{S23}$$

  – Switch from refractory state to growing state with probability $p = 1 - \exp(-1/T_r)$:

$$\begin{aligned} &s^{(\alpha)}(t) = 1 \\ &\Rightarrow s^{(\alpha)}(t+1) = \begin{cases} 2 & (p), \\ 1 & (1 - p), \end{cases} \end{aligned} \tag{S24}$$

  where the terms in the brackets denote the respective probability.

  – Exponential growth in interphase over a period of $T_g$:

$$\begin{aligned} &s^{(\alpha)}(t) = 2 \\ &\Rightarrow \kappa_A^{(\alpha)}(t+1) = \kappa_A^{(\alpha)}(t) \cdot 0.5^{1/T_g} \\ &\Rightarrow \kappa_P^{(\alpha)}(t+1) = \kappa_P^{(\alpha)}(t) \cdot 0.25^{1/T_g} \end{aligned} \tag{S25}$$

  – Switch from interphase to mitosis:

$$\begin{aligned} &s^{(\alpha)}(\tau) = 2 \text{ for all } \tau \in [t - T_g, t] \\ &\Rightarrow s^{(\alpha)}(t + 1) = 3. \end{aligned} \tag{S26}$$

  During cell division, cell motility is switched off and the cytoskeletal density relaxes to the neutral state according to Eq. 3c.

  – Perform cell division, reset area and perimeter stiffness and exit mitotic phase:

$$\begin{aligned} &s^{(\alpha)}(\tau) = 3 \text{ for all } \tau \in [t - T_m, t] \\ &\Rightarrow \text{ divide cell } \alpha \text{ into cells } (\alpha, \beta) \\ &\Rightarrow \kappa_A^{(\alpha,\beta)}(t+1) = \kappa_{A,0} \\ &\Rightarrow \kappa_P^{(\alpha,\beta)}(t+1) = \kappa_{P,0} \\ &\Rightarrow s^{(\alpha,\beta)}(t + 1) = 0. \end{aligned} \tag{S27}$$

  – Cell motility is restored after cell division.

  – If none of the above rules applies do not perform any changes.

### G. Numerical computation of stress in a tissue

In the section describing the numerical results on tissue expansion, the stress distribution in the tissue is shown in the kymographs Figs. 6(b,e). Hereafter we explain how the stress tensor for each cell in the tissue can be computed from the forces acting on the cell's membrane pixels in the Monte Carlo simulation. The mean value of the stress tensor in a deformed body can be calculated numerically from the formula

$$\bar{\sigma}_{ij}^{(\alpha)} = \frac{\ell}{2A^{(\alpha)}} \sum_{\mathbf{x}_k \in \mathcal{B}^{(\alpha)}} \left( f_k^i x_k'^j + f_k^j x_k'^i \right) , \tag{S28}$$

which is a discretized version of the surface integral in Ref. [21]. Here $\mathbf{f_k}$ is the force acting on the membrane element $\mathbf{x}_k$ of cell $\alpha$, $\mathbf{x'_k} = \mathbf{x}_k - \mathbf{x}_{\mathrm{com}}^{(\alpha)}$ is the position of the element with respect to the center of mass $\mathbf{x}_{\mathrm{com}}^{(\alpha)}$ of the cell, and the superscripts $i$ and $j$ are Cartesian indices. The forces $\mathbf{f_k}$ can be computed from the energy differences of all possible protrusion and retraction events originating from $\mathbf{x}_k$,

$$\begin{aligned} \mathbf{f}_k = &- \sum_{\mathbf{x}_l \in \mathcal{N}_k} \frac{\Delta\mathcal{H}(\mathcal{T}_{\mathrm{pro}})}{\|\mathbf{x}_l - \mathbf{x}_k\|} \frac{\mathbf{x}_l - \mathbf{x}_k}{\|\mathbf{x}_l - \mathbf{x}_k\|} \\ &- {\sum_{\mathbf{x}_l \in \mathcal{N}_k}}' \frac{\Delta\mathcal{H}(\mathcal{T}_{\mathrm{ret}})}{\|\mathbf{x}_k - \mathbf{x}_l\|} \frac{\mathbf{x}_k - \mathbf{x}_l}{\|\mathbf{x}_k - \mathbf{x}_l\|} , \end{aligned} \tag{S29}$$

where the prime indicates a sum over substrate pixels only, i.e. pixels with $c(\mathbf{x}_l) < 0$, and where $\Delta\mathcal{H} \equiv \mathcal{H}_{\mathrm{cont}} + \mathcal{H}_{\mathrm{adh}} + \mathcal{H}_{\mathrm{cyto}}$.

#### H. Numerical computation of the cell shape

We use two complementary measures for the cell shape. The first is a simple measure for the deviation of an object from a circle (extension):

$$K = 1 - \frac{4\pi A}{P^2} . \tag{S30}$$

It becomes zero if the object is a circle and becomes 1 if the object is a line. The second measure for the cell shape is obtained from a principle components analysis of the cell shape. Specifically, we compute the covariance matrix of the point cloud representing the cell $\mathcal{D}^{(\alpha)}$:

$$\mathrm{Cov}(\mathcal{D}^{(\alpha)}) = \begin{pmatrix} A_{XX} & A_{XY} \\ A_{XY} & A_{YY} \end{pmatrix} \tag{S31}$$

and then compute the corresponding ratio of the larger to the smaller eigenvalue $l_{\mathrm{long}}/l_{\mathrm{short}}$ (aspect ratio). In coordinates relative to the cell center of mass $\tilde{\mathbf{x}}_i = (\tilde{x}_i, \tilde{y}_i) = \mathbf{x}_i - \mathbf{x}_{\mathrm{com}}^{(\alpha)}$, the components of the covariance matrix are given by

$$A_{XX} = \frac{\sum_{\mathbf{x}_i \in \mathcal{D}^{(\alpha)}} \tilde{x}_i \, \tilde{x}_i}{\sum_{\mathbf{x}_i \in \mathcal{C}} 1} , \tag{S32}$$

$$A_{XY} = \frac{\sum_{\mathbf{x}_i \in \mathcal{D}^{(\alpha)}} \tilde{x}_i \, \tilde{y}_i}{\sum_{\mathbf{x}_i \in \mathcal{C}} 1} , \tag{S33}$$

$$A_{YY} = \frac{\sum_{\mathbf{x}_i \in \mathcal{D}^{(\alpha)}} \tilde{y}_i \, \tilde{y}_i}{\sum_{\mathbf{x}_i \in \mathcal{C}} 1} . \tag{S34}$$

### S.II. PARAMETER SCREENING IN SILICO

In this section we provide additional analysis of the model parameters beyond what is already shown in the main text.

We explore how the motility and morphology of single cells depend on key model parameters like the polarizability range $\Delta Q$, the perimeter stiffness $\kappa_P$, the signaling radius $R$, and dissipative effects $D$ in the interaction between the cell and the substrate. Here, we find that single cells elongate and become more motile (both higher cell speed and persistence time of directed migration) with increasing specific polarizability $\Delta Q/\kappa_P$ With increasing cell-substrate dissipation, cell migration is inhibited and the cells round up. Furthermore, we can map the migratory behavior of the cell on their shape and conclude that elongated cells have higher motility (persitence and speed). The effects of increasing the signaling radius are more complex: while for low signaling radii an increase of the signaling radius leads to higher coordination of cytoskeletal activity in the leading edge of the cell, for high signaling radii this effect is reversed as protrusion/retraction events at one edge of the cell begin to affect the opposing edge. Using the signaling radius as a control parameter, we can switch between amoeboidal and persistent cell migration.

Furthermore, we explore all three rotational phases $\mathcal{R}_1$, $\mathcal{R}_2$ and $\mathcal{R}_3$ within confinements of varying size and constant cell density. In the $\mathcal{R}_1$-phase, the cell clusters rotate slowly and reorient frequently their direction of rotation. With increasing cell count, macroscopic rotation of the cluster stops. In the highly coordinated $\mathcal{R}_2$ and $\mathcal{R}_3$-phases, we find scale-free behavior such that there is always a macroscopic rotation of the whole cell population regardless of the cell count and corresponding container size.

We also explore the parameter space of tissue-simulations. There, we find that an increased cell-cell dissipation $\Delta B$ impairs monolayer growth, while at the same time increasing the front roughness. Similarly, an increased cell-substrate dissipation $D$ also impairs monolayer growth. In contrast, increasing the maximal cell polarity $\Delta Q$ improves monolayer growth and also increases the front roughness. We thus find that the speed of monolayer expansion depends on whether it is dominated by cell migration or cell proliferation, with the former improving growth by a better exploration of the cell-free area.

#### A. Single cell size

To rationalize our choice of the cell growth algorithm [Sec. S.I F], we have explored the shape and motility of differently sized cells. To this end, we have varied the area stiffness parameter $\kappa_A$ for different values of perimeter stiffness $\kappa_P$, while keeping all other parameters constant. We find that the area occupied by the motile cell

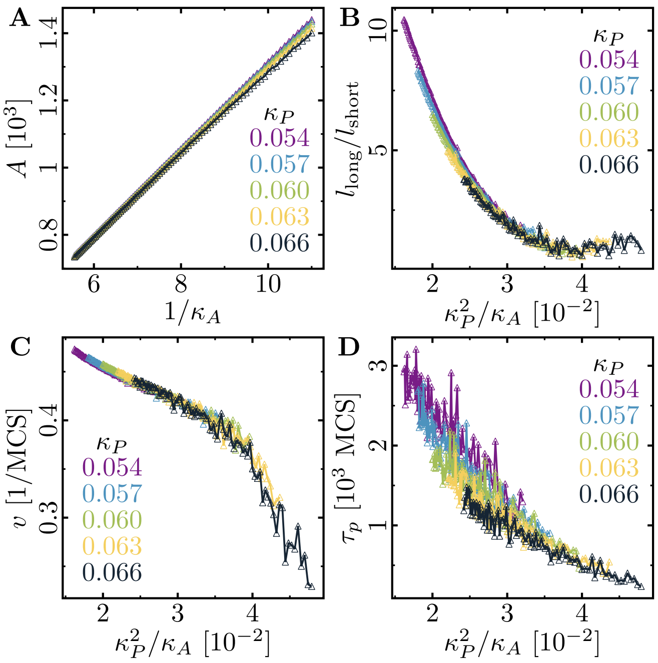


FIG. S4. **Role of area stiffness $\kappa_A$ for cell size and motility.** **(A)** The cell area increases linearly with $1/\kappa_A$. The aspect ratio **(B)**, speed **(C)** and persistence **(D)** of the cell decrease with increasing cell size. In the simulations, the area elasticity was varied in the interval $\kappa_A \in [0.09\ldots0.18]$, and the membrane elasticity was chosen from $\kappa_P \in \{0.054, 0.057, 0.060, 0.063, 0.066\}$. Fixed parameters: average cytoskeletal density $(Q+q)/2 = 225$; maximal cell polarity $\Delta Q = 50$; signaling radius $R = 5$; cytoskeletal update rate $\mu = 0.1$; cell-substrate dissipation $D = 0$; cell-substrate adhesion penalty $\varphi = 0$.

increases linearly with $1/\kappa_A$ [Fig. S4A]. In particular, the cell area can be approximated quite well by the area of an immotile and equilibrated cell with uniform $\rho = (q+Q)/2$ (fit not shown):

$$A = \frac{q + Q - 4\pi\sqrt{3}\kappa_P}{2\sqrt{3}\kappa_A} \propto 1/\kappa_A\,. \tag{S35}$$

Furthermore, we find that with increasing size, and all other parameters constant, cells become rounder, slower, and less persistent [Fig. S4B-D]. To intuitively explain this phenomenology, let us compare a cell of size $A_{\mathrm{ref}}$ with a cell of size $c\,A_{\mathrm{ref}}$, where $c \in [1\ldots2]$, with the respective area stiffnesses $\kappa_A$ and $\kappa_A/c$. While the smaller cell has a perimeter $P_{\mathrm{ref}}$, neglecting geometric effects the larger cell has a larger perimeter $\approx \sqrt{c}P_{\mathrm{ref}}$. Hence, the larger cell has to pay a larger energy cost (roughly by a factor $\sqrt{c}$) to ruffle its membrane by adding segments and therefore increasing its perimeter. Meanwhile, both the energy gain from the cytoskeletal density $\rho$ and the energy cost for increasing the cell area (as $A \cdot \kappa_A = \text{cst.}$) are the same for both cells. Due to the increased energy cost for adding membrane segments, larger cells finds find it more difficult to polarize, and are therefore rounder, slower and less persistent.

To offset this increased energy cost for adding membrane segments to the cell, we can scale the perimeter stiffness of the larger cell to $\kappa_P/\sqrt{c}$, such that $P \cdot \kappa_P \approx$ cst.. We would then predict that the ratio $\kappa_P^2/\kappa_A$ is constant for differently sized cells of similar shape, speed and persistence time. The same relation can also be obtained by realizing that different amounts of hexagons occupied by two otherwise identical cells in terms of their corresponding Hamiltonian simply stem from a different *discretization* of said cells, controlled by the parameter $\kappa_A$. Interestingly, we observe such a data collapse for the aspect ratio $l_{\text{long}}/l_{\text{short}}$ and the velocity $v$ of the cells onto two respective master curves depending on the ratio $\kappa_P^2/\kappa_A$ [Fig. S4B,C]. While the proposed data collapse for the persistence time of the cell [Fig. S4D] is somewhat unsatisfactory, this may be owed to the following effect: by keeping $R$ constant we have actually varied the ratio of the area that the signaling molecules typically explore due to diffusion and the area of the cell $R^2/A$. We speculate that all observed quantities collapse unto respective master curves $f(\Delta Q\sqrt{\kappa_A}/\kappa_P) \cdot g(R\sqrt{\kappa_A})$.

#### B. Single cell shape and dynamics

To learn more how the observed morphology and motility phenotypes depend on the model parameters, we studied how the aspect ratio $l_{\text{long}}/l_{\text{short}}$ obtained from a principal components analysis of the point cloud representing the cell, the elongation $1 - 4\pi A/P^2$, the persistence time $\tau_p$ and the the cell speed $v$ depend on the following parameters: polarizability range $\Delta Q$, rigidity parameter $\kappa_P$ for the cell perimeter, and the signaling radius $R$ [Fig. S5, S7].

As can be inferred from Fig. S5, there is data collapse for all three observables. They do not depend on the polarizability range $\Delta Q$ and the perimeter stiffness $\kappa_P$ separately but only on their ratio $\Delta Q/\kappa_P$. Heuristically this may be explained as follows: Let us consider cells of identical size or discretization, and therefore fixed area elasticity $\kappa_A$. Then, the perimeter stiffness term in the Hamiltonian will punish an increase in cellular perimeter, and intuitively $P \propto 1/\kappa_P$. To illustrate this, let us look at a simple extreme: for $\kappa_P \equiv 0$, an extremely elongated cell with an infinitesimal width $W \to 0$ and infinite length $L \to \infty$ such that the total area $A = W\,L$ remains fixed would not be energetically penalized compared to a round cell. Let us take a step further and consider such a profoundly elongated cell, where the elongation of the cell is facilitated by the polarization mechanism driving the cell out of equilibrium. For such an elongated cell, the width of the cell $W$ takes up most of the cell perimeter $W \sim P/2$. If such a cell moves an infinitesimal distance $\delta\mathbf{x}$, while overall preserving its shape (both area and perimeter) and cytoskeletal density, the energy difference driving this motion is given by

$$\delta\mathcal{H}_P = \delta \int_A d^2\mathbf{x}\,\rho(\mathbf{x}) = \delta\mathbf{x} \int_A d^2\mathbf{x}\,\nabla\rho(\mathbf{x})\,. \qquad \text{(S36)}$$

A crude estimation then yields for elongated cells:

$$\delta\mathcal{H}_P \approx |\delta\mathbf{x}|\,\frac{\Delta Q}{L} A = |\delta\mathbf{x}|\,\Delta Q W \ . \qquad \text{(S37)}$$

In this estimation we have sweeped most of the dynamics under the carpet, including the whole polarization mechanism. Nevertheless, we can still make several statements on the model behavior in the parameter limits: For elongated cells $\Delta Q \to \infty$ or $\kappa_P \to 0$, the energy difference driving migration will also diverge $\delta\mathcal{H}_P \to \infty$ [Eq. S37]. For round cells $\Delta Q \to 0$ or $\kappa_P \to \infty$, our estimate does not hold. However, the energy difference driving migration will still vanish $\delta\mathcal{H}_P \to 0$: because of the vanishing energy gain or infinite energy penalty for ruffling the cell membrane, respectively, the cell will equilibrate to a constant cytoskeletal density $\nabla\rho \to 0$ [Eq. S36]. The

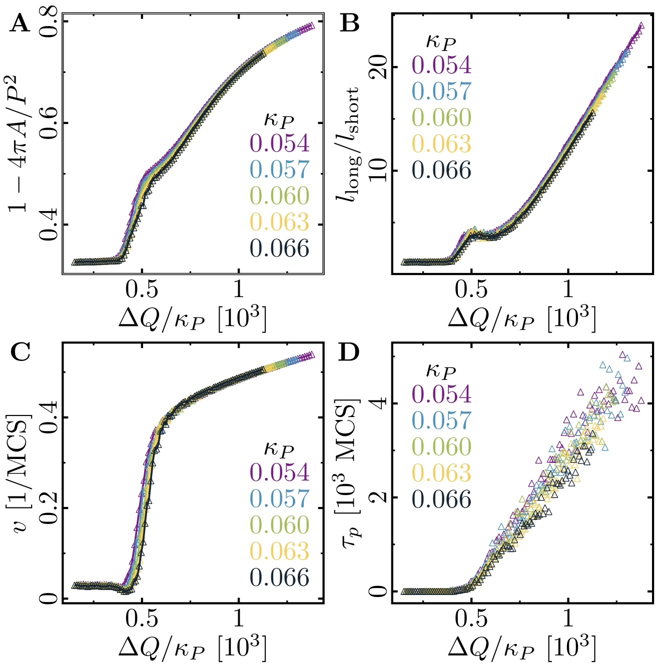


FIG. S5. **Parameter study of morphology and motility phenotypes depending on the polarizability $\Delta Q$.** The aspect ratio $l_{\text{long}}/l_{\text{short}}$ (A), the extension $1 - 4\pi A/P^2$ (B), the persistence time $\tau_p$ (C), and the speed $v$ (D) as a function of the ratio between polarizability range $\Delta Q$ and rigidity parameter for the cell periphery $\kappa_P$ for a set of values for $\kappa_P$ indicated in the graphs. In the simulations, the maximal cell polarity was varied in the interval $\Delta Q \in [10 \dots 75]$, and the membrane elasticity was chosen from $\kappa_P \in \{0.054, 0.057, 0.060, 0.063, 0.066\}$. Fixed parameters: average cytoskeletal density $(Q+q)/2 = 225$; area elasticity $\kappa_A = 0.18$; signaling radius $R = 5$; cytoskeletal update rate $\mu = 0.1$; cell-substrate dissipation $D = 0$; cell-substrate adhesion penalty $\varphi = 0$.

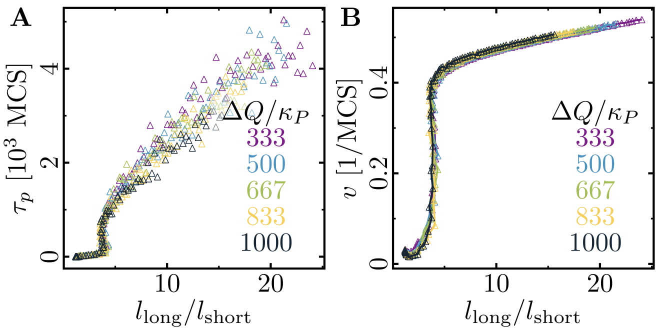


FIG. S6. **Replotted values of Fig. S5.** Cell speed and cell persistence are correlated with the aspect ratio of the cell.

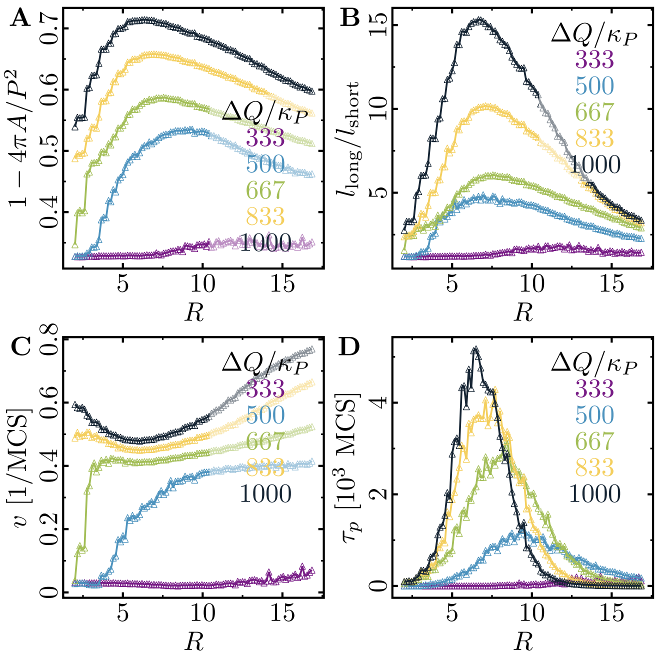


FIG. S7. **Parameter study of morphology and motility phenotypes depending on the signaling radius $R$.** The aspect ratio $l_{\mathrm{long}}/l_{\mathrm{short}}$ (A), the extension $1-4\pi A/P^2$ (B), the persistence time $\tau_p$ (C), and the speed $v$ (D) as a function of the signaling radius $R$ for a set of values for $\kappa_P$ indicated in the graphs. In the simulations, the signaling radius was varied in the interval $R \in [1\ldots17]$, and the maximal cell polarity $\Delta Q \in \{20, 30, 40, 50, 60\}$. Fixed parameters: average cytoskeletal density $(Q+q)/2 = 225$; area elasticity $\kappa_A = 0.18$; cytoskeletal update rate $\mu = 0.1$; membrane elasticity $\kappa_P = 0.060$; cell-substrate dissipation $D = 0$; cell-substrate adhesion penalty $\varphi = 0$.

simplest hypothesis that is compatible with the model behavior in the parameter limits is then $\delta\mathcal{H} \sim \Delta Q/\kappa_P$.

Note that because of this universal dependence of all the mentioned quantities on the ratio $\Delta Q/\kappa_P$, there is a correlation between cell morphology (aspect ratio) and cell motility (speed and persistence time). In particular, there are direct relations of the persistence time and the speed with the aspect ratio [Fig. S6]. This correlation between morphology and motility was previously also observed in experiments on Fish Keratocytes [22].

As a function of the scaling variable $\Delta Q/\kappa_P$, there is interesting behavior. We observe that the aspect ratio, the extension, the persistence time $\tau_p$ and the speed of the cell all increase with $\Delta Q/\kappa_P$ [Fig. S5]. In particular, we note that there is a lower threshold of $\Delta Q/\kappa_P \sim 400$, below which cells are round and immotile [Fig. S5A,D]. Above an upper threshold $\Delta Q/\kappa_P \sim 750$ the cells show a linear increase with $\Delta Q/\kappa_P$ for their aspect ratio, their speed, and their persistence time [Fig. S5A-C]. To illustrate a correlation between cell shape and cell migration, we replot the values of Fig. S5 and observe an overall increase in both cell persistence and cell speed with its aspect ratio [Fig. S6].

We also explored the role of the signaling radius $R$ [Fig. S7D], and made the following observations: Depending on the cell polarizability ($\Delta Q$), there is an optimal signaling radius that shows both maximal cell elongation and maximal cell persistence [Fig. S7A,C]. We understand this as an effect due to the ratio of the area that the signaling molecules typically explore due to diffusion ($R^2$) and the total area and shape of the cell. As the signaling radius defines the range in which the signaling molecules typically diffuse during a single Monte-Carlo step and cellular events update the cell polarization, it is the essential parameter for the formation of stable polarization fronts. However, if $R$ becomes too large, then the retraction events at the retracting edge of the cell also start to influence the leading edge and vice versa. This will lead to an overall decrease in the effective polarization of the cell ($\Delta Q_{eff} \leq \Delta Q$) and thus lead to a rounding of the cell (Compare Fig. S5A).

Cells with a low polarizability ($\Delta Q$) need a large signaling radius to feed the positive feedback mechanism and form a single large cell front. In contrast, highly polarizable cells can already sustain the positive feedback mechanism with a short signaling radius and easily form at least one (or even multiple competing) short cell front(s). With increasing signaling radius, these cell fronts become increasingly correlated and finally merge. Surprisingly, we observe for highly polarizable cells that the cell speed initially decreases with increasing signaling radius [Fig. S7C], in contrast to the behavior observed in cells with low polarizability. Furthermore, we there observe an increase in cell speed for high signaling radii, although cell persistence has dropped to small values [Fig. S7C,D]. To find an intuitive explanation for these observations, we inspected time-lapse videos of the cell at high polarizability ($\Delta Q/\kappa_P = 1000$; cf. Supplemental Videos 1–3), which show a a qualitative shift in cell behavior: For $R = 2$ (Supplemental Video 1), short polarization fronts 'pull' the cell behind them, allowing for transient polarization and movement along the long axis of the cell. For $R = 6$ (Supplemental Video 2), broad and correlated polarization fronts emerge, and both the cell polarization and movement always orient themselves along the short axis of the cell. Let us illustrate why

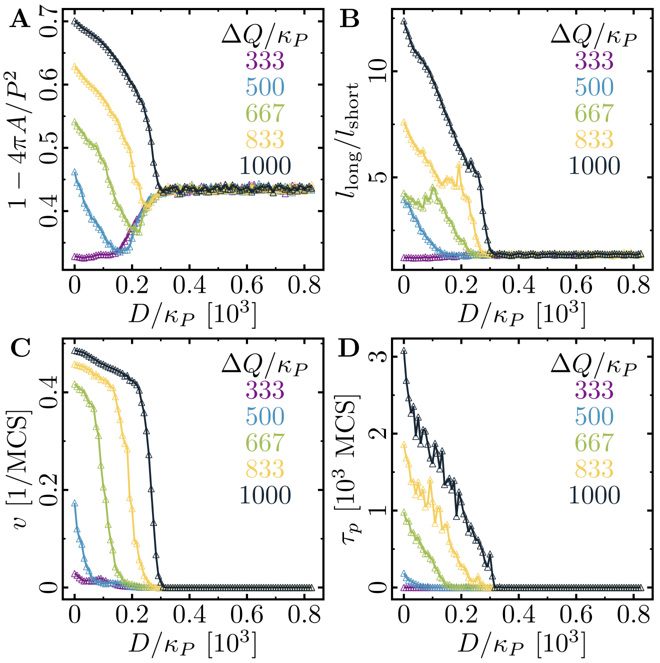


FIG. S8. **Role of substrate dissipation for cells of varying polarity $\Delta Q$.** Cell speed $v$ (A), persistence time $\tau_p$ (B), and aspect ratio (C) as a function of substrate dissipation $D$ for a series of values for the perimeter stiffness $\kappa_P$ indicated in the graphs. In the simulations, the substrate dissipation was varied in the interval $D \in [0 \dots 50]$, and the maximal cell polarity $\Delta Q \in \{20, 30, 40, 50, 60\}$. Fixed parameters: average cytoskeletal density $(Q+q)/2 = 225$; area elasticity $\kappa_A = 0.18$; membrane elasticity $\kappa_P = 0.060$; cytoskeletal update rate $\mu = 0.1$; cell-substrate dissipation $D = 0$; cell-substrate adhesion penalty $\varphi = 0$.

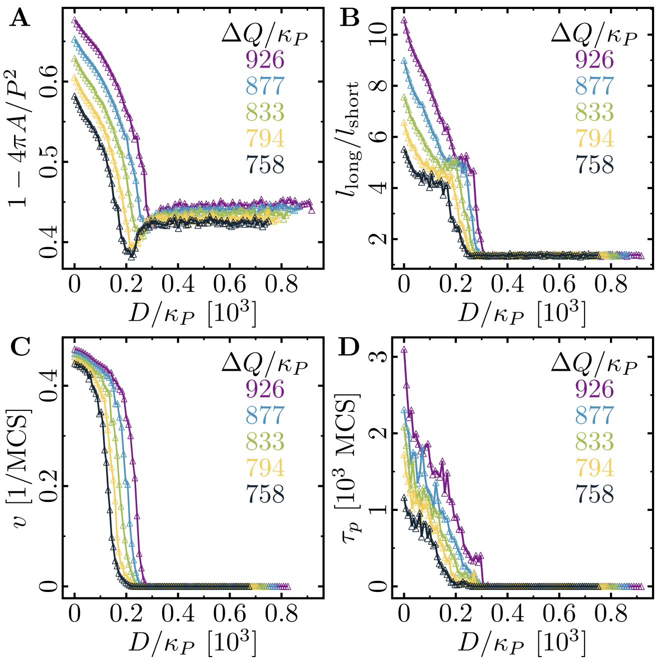


FIG. S9. **Role of substrate dissipation for cells of varying membrane stiffness $\kappa_P$.** Cell speed $v$ (A), persistence time $\tau_p$ (B), and aspect ratio (C) as a function of substrate dissipation $D$ for a series of values for the perimeter stiffness $\kappa_P$ indicated in the graphs. In the simulations, the substrate dissipation was varied in the interval $D \in [0 \dots 50]$, and the membrane elasticity $\kappa_P \in \{0.054, 0.057, 0.060, 0.063, 0.066\}$. Fixed parameters: average cytoskeletal density $(Q+q)/2 = 225$; area elasticity $\kappa_A = 0.18$; maximal cell polarity $\Delta Q = 50$; cytoskeletal update rate $\mu = 0.1$; cell-substrate dissipation $D = 0$; cell-substrate adhesion penalty $\varphi = 0$.

motion along the long axis is faster in our simulations. Consider two equally sized cells with widths $W_\perp$ and $W_\parallel$ ($W_\perp > W_\parallel$) that move perpendicular or parallel to their long axis, respectivly. Furthermore, each cell has a total budget $M$ of move attempts during a MCS. If we assume that only move attempts in the respective direction of motion are successful, then the cell that moves parallel to its long axis will travel at a speed $v_0 M/W_\parallel$, while the other cell will only travel at a slower speed $v_0 M/W_\perp < v_0 M/W_\parallel$. Here, $v_0\, M = 0.5a\, M/\mathrm{MCS} = \Delta A/\Delta t$ corresponds to the cell area gained at the leading edge and lost at the trailing edge per MCS. For $R = 15$ (Supplemental Video 3), we observe circular motion of the cell; because of the high signaling radius, signals originating from the trailing edge affect the leading edge of the cell and vice versa. Due to this circular motion, the cell exhibits a non-zero speed and a vanishing persistence time of directed migration.

Finally, we have also studied the effect of cell-substrate dissipation (see Sec. S.I D 5) on cell morphology and motility. We have varied $D$ for different values of $\Delta Q$ and $\kappa_P$; however, we were not able to achieve a data collapse in $D$ [Figs. S8,S9]. We observe that with increasing cell-substrate dissipation, cells become round and cease migrating. This can be illustrated as follows: Consider a situation where the cell conquers a new hexagon at its prospective leading edge. Because the cell on average tends to constrain its area and perimeter while migrating, it needs to lose a different hexagon at its prospective trailing edge. However, this retraction at the trailing edge is energetically penalized and thus cell displacement from its initial position and the positive feedback leading to cell polarization are effectively inhibited. With increasing cell-substrate dissipation, retraction events are further penalized and the cell 'sticks' to the substrate at its trailing edge, preventing persistent motion of the cell. Additionally, to further illustrate the correlation between cell shape and cell migration, we have replotted the values of Fig. S8 [Fig. S10].

#### C. Cells in circular confinement

In this section we report on additional parameter studies of the dynamics of cells enclosed in a circular confinement [Figs. S11, S12 and S13]. Specifically, we investigate

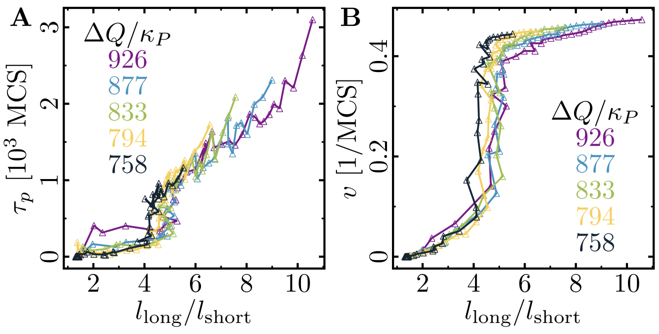


FIG. S10. **Replotted values of Fig. S9.** Cell speed and cell persistence are correlated with the aspect ratio of the cell (cf. Fig. S6).

how the radius of circular confinement affects the synchronized rotation of the cell population. We performed simulations with a densely populated circular confinement and varied the confinement radius, while keeping the cell density constant. The parameters are chosen such that a population consisting of 4 cells (cf. main text, Fig. 4) would rotate in the $\mathcal{R}_1$, $\mathcal{R}_2$ or $\mathcal{R}_3$-phase, respectively. We studied mean angular velocity

$$\omega := \hat{\mathbf{e}}_z \left\langle \frac{\mathbf{v} \times \mathbf{r}}{r^2} \right\rangle , \qquad \text{(S38)}$$

averaged over all cells in confinement, where $\mathbf{v}$ and $r$ are the velocity relative to and distance to the center of mass of the cell cluster, respectively.

In the lowly polarizable $\mathcal{R}_1$-phase, small cell populations rotate in a highly synchronized way, and rotation is maximized for populations of 7 cells per confinement [Fig. S11A]. As can be inferred from the time traces and snapshots [Fig. S11B,C], cells at a given time all synchronously move in the same direction and randomly switch between clockwise and counter-clockwise rotation; the switching rate decreases with increasing size of the cell population. Upon increasing the cell count and concomitantly the confinement size, global rotation of the cell population gradually vanishes [Fig. S11A].

Unlike in the $\mathcal{R}_1$-phase, we observe that in the highly polarizable $\mathcal{R}_2$ and $\mathcal{R}_3$-phases populations of all sizes rotate in a highly synchronized way [Figs. S12A and S13A]. There, the dependence of $\langle|\omega|\rangle$ on the populations size $N$ can be fitted by a power law of the form $\langle|\omega|\rangle \propto N^{-1/2} \propto r_0^{-1}$. This inverse proportionality of average angular velocity on the confinement size $r_0$ implies total rotational order, with every cell moving at a constant velocity $|\mathbf{v}_{\rm rot}| \approx 0.08$ ($\mathbf{v}_{\rm rot} \perp \mathbf{r}$). Furthermore, in the $\mathcal{R}_2$, and $\mathcal{R}_3$-phases we have observed only scarcely switching of the rotational direction of cell clusters; e.g. for 4-cell clusters in the $\mathcal{R}_3$-phase.

Interestingly, fluctuations in the angular velocity $\sigma_\omega$ change in a highly non-monotonic fashion with the cell count and concomitantly the confinement size. Certain cell counts exhibit especially high fluctuations of

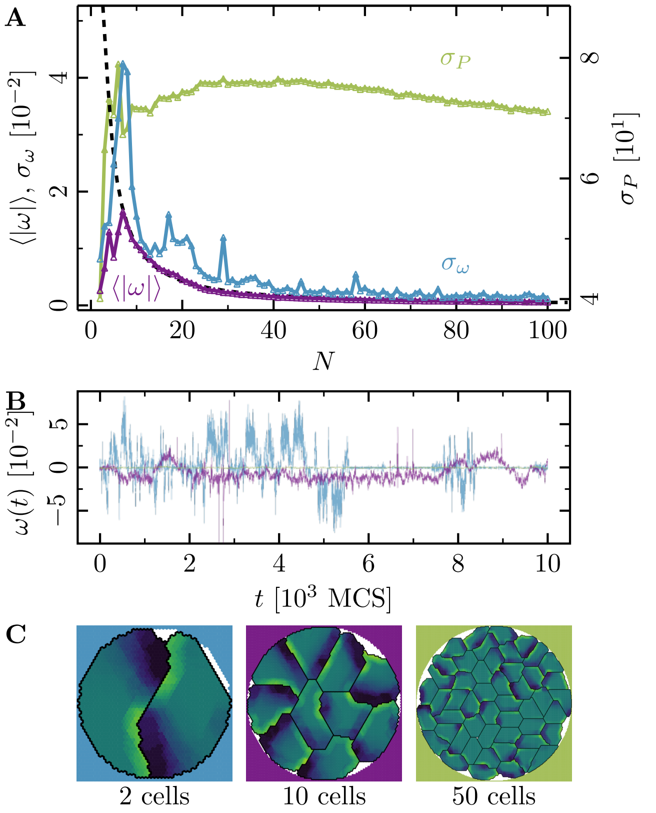


FIG. S11. **Collective motion for varying number of cells.** ($N$ cell systems, confinement radius $r_0 = \sqrt{234\,N}$; area stiffness $\kappa_A = 0.18$; average cytoskeletal density $(Q+q)/2 = 225$; maximal cell polarity $\Delta Q = 28$; signaling radius $R = 5$; cytoskeletal update rate $\mu = 0.1$; cell-cell adhesion $B = 0$; cell-cell dissipation $\Delta B = 7$; cell-substrate dissipation $D = 0$; cell-substrate adhesion penalty $\varphi = 0$ ($r < r_0$), $\varphi \to -\infty$ ($r > r_0$)). For this choice of parameters 4-cell populations rotate in the $\mathcal{R}_1$-phase. We observe a similar behavior here: the cell clusters rotate slowly and reorient frequently. **(A)** Characteristic observables of collective cell rotation at different values of the cell perimeter stiffness parameter $\kappa_P$: mean ($\langle\omega\rangle$) and standard deviation ($\sigma_\omega$) of angular velocity of cell motion, and the standard deviation of the cell shape variability ($\sigma_P$). The black line corresponds to a power-law fit of the form $\langle|\omega|\rangle \propto N^{-k/2} \propto r_0^{-k}$ with the fitted exponent $k \approx 8/3$. **(B)** Representative angular trajectories and **(C)** cell shapes (color code represents cell polarization; cf. Fig. 1) for the different parameter regimes as described in the main text.

the mean angular velocity (e.g. 5 cells in the $\mathcal{R}_1$-phase [Fig. S11A]; 3 or 10 cells in the $\mathcal{R}_2$-phase [Fig. S12A]; 3 cells in the $\mathcal{R}_3$-phase Fig. S13A), which can likely be attributed to frustration of the cells in the population center [cf. $N = 10$ cells in Fig. S12C].

#### D. Velocity and roughness of spreading tissue

We have studied the velocity and roughness of spreading tissue, while varying cell-cell dissipation $\Delta B$, cell-

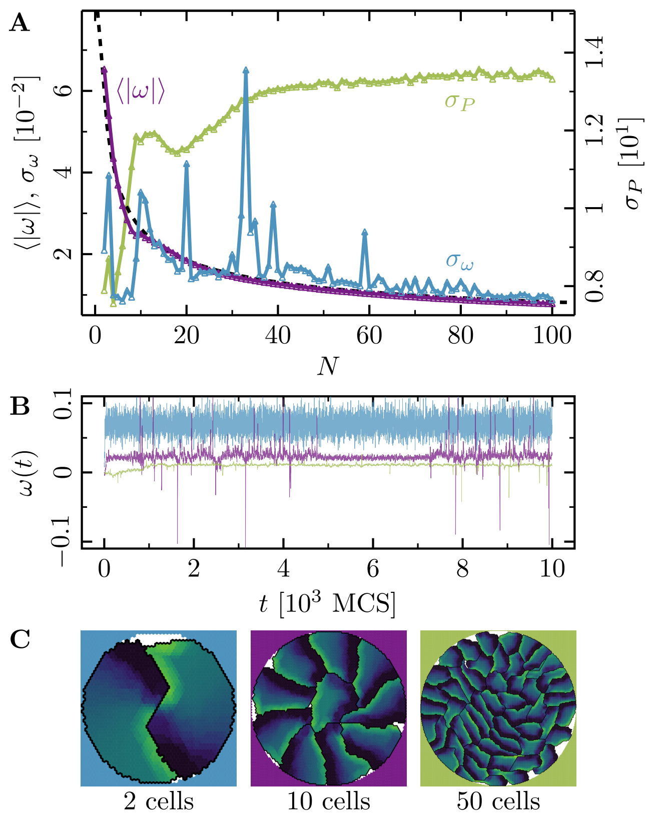


FIG. S12. **Collective motion for varying number of cells.** ($N$ cell systems, confinement radius $r_0 = \sqrt{234\,N}$; area stiffness $\kappa_A = 0.18$; average cytoskeletal density $(Q+q)/2 = 225$; maximal cell polarity $\Delta Q = 50$; signaling radius $R = 5$; cytoskeletal update rate $\mu = 0.1$; cell-cell adhesion $B = 0$; cell-cell dissipation $\Delta B = 7$; cell-substrate dissipation $D = 0$; cell-substrate adhesion penalty $\varphi = 0$ ($r < r_0$), $\varphi \to -\infty$ ($r > r_0$)). For this choice of parameters 4-cell populations rotate in the $\mathcal{R}_2$-phase. We observe a similar behavior here: highly correlated rotations with no changes in rotational direction. **(A)** Characteristic observables of collective cell rotation at different values of the cell perimeter stiffness parameter $\kappa_P$: mean ($\langle\omega\rangle$) and standard deviation ($\sigma_\omega$) of angular velocity of cell motion, and the standard deviation of the cell shape variability ($\sigma_P$). The black line corresponds to a power-law fit of the form $\langle|\omega|\rangle \propto N^{-1/2} \propto r_0^{-1}$. **(B)** Representative angular trajectories and **(C)** cell shapes (color code represents cell polarization; cf. Fig. 1) for the different parameter regimes as described in the main text.

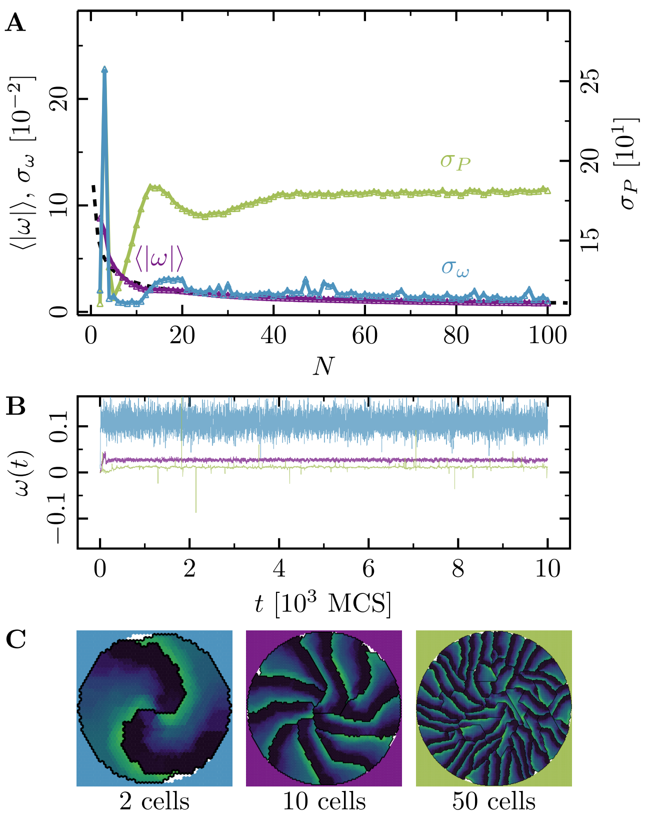


FIG. S13. **Collective motion for varying number of cells.** ($N$ cell systems, confinement radius $r_0 = \sqrt{234\,N}$; area stiffness $\kappa_A = 0.18$; average cytoskeletal density $(Q+q)/2 = 225$; maximal cell polarity $\Delta Q = 70$; signaling radius $R = 5$; cytoskeletal update rate $\mu = 0.1$; cell-cell adhesion $B = 0$; cell-cell dissipation $\Delta B = 7$; cell-substrate dissipation $D = 0$; cell-substrate adhesion penalty $\varphi = 0$ ($r < r_0$), $\varphi \to -\infty$ ($r > r_0$)). For this choice of parameters 4-cell populations rotate in the $\mathcal{R}_3$-phase. We observe a similar behavior here: highly correlated rotations. **(A)** Characteristic observables of collective cell rotation at different values of the cell perimeter stiffness parameter $\kappa_P$: mean ($\langle\omega\rangle$) and standard deviation ($\sigma_\omega$) of angular velocity of cell motion, and the standard deviation of the cell shape variability ($\sigma_P$). The black line corresponds to a power-law fit of the form $\langle|\omega|\rangle \propto N^{-1/2} \propto r_0^{-1}$. **(B)** Representative angular trajectories and **(C)** cell shapes (color code represents cell polarization; cf. Fig. 1) for the different parameter regimes as described in the main text.

substrate dissipation $D$ and cell polarizability $\Delta Q$.

First, let us introduce the observables that we're interested in. Let $\mathcal{X}_{>/<}$ be the sets of $x$-coordinates of the left and right outermost edges of the cell sheet. Our *in silico* setup is axially symmetric with respect to the $y$-axis. This initial symmetry persists, as the cell fronts advance towards the cell-free area with the same average speed. Hence, it is not needed to consider the two cell fronts separately, and we can instead consider the set of *unsigned* front positions $\mathcal{X} := \text{abs}(\mathcal{X}_{>/<})$. Then, we define the average front position as $x_\text{F} := \mathbb{E}(\mathcal{X})$ and the front roughness as $\sigma_\text{F}^2 := \text{Var}(\mathcal{X})$. In particular, we study the total growth of the tissue over the course of 500 MCS, which is captured by the maximal position of the front $\max(x_\text{F})$, as well as the maximal roughness of the front $\max(\sigma_\text{F})$. We have chosen our parameters such that a cell takes a total amount of 200 MCS to divide, provided that it exceeds the threshold size of a solitary reference cell $A_\text{ref}$. Because the first daughter cells may only appear after 200 MCS have passed, we exclude this initial period from the measurements of the maximal front position and roughness, respectively. Additionally, we provide some exemplary time traces of the front evolution.

Our simulations show that the cell sheet expands slower with increasing cell-cell dissipation $\Delta B$ [Fig. S14A,B], because the dissipation penalizes cells sliding past each other. At the same time, the cell sheet also becomes slightly rougher with increasing cell-cell dissipation $\Delta B$ [Fig. S14A].

We also investigate how the monolayer expansion and front roughness depend on cell-substrate dissipation, $D$ [Fig. S15]. Before we turn to the monolayer, let us recall the observed single-cell behavior in the previous section [Sec. S.II B]: for high enough cell-substrate dissipation $D$ (typically of the same order of magnitude as the maximal cell polarity $\Delta Q$) cell migration is switched off [Figs. S8 and S8]. Extrapolating the single-cell results, we expect that the same holds also for collectives of cells and that cell migration does not play a role for high cell-substrate dissipation. Indeed, with increasing cell-substrate dissipation, the monolayer expands slower, until this effect appears to saturate at a threshold value $D^\star \approx 5$ [Fig. S15A,B]. Following this line of argument, monolayer growth is slowed down if we suppress cell migration and thus move the cell monolayer towards a proliferation-dominated mode of expansion.

What about the inverse? Is the monolayer growth increased if we enhance cell migration and thus move the cell monolayer towards a migration-dominated mode of expansion? To test this hypothesis, we have analyzed how the monolayer growth and front roughness depend on the maximal cell polarity $\Delta Q$. As predicted, monolayer growth increases with the maximal cell polarity $\Delta Q$ [Fig. S16A,B], because an increased amount cells exceed the threshold size to switch to mitosis (cf. the stretching of bulk cells in the monolayer in Fig. 5B). Additionally, we also find that the front roughness increases with increasing polarizability $\Delta Q$.

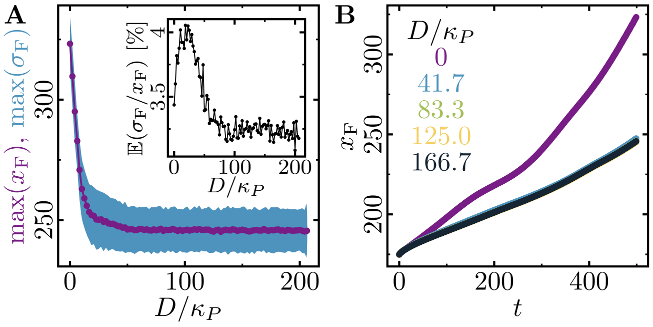


FIG. S15. Cell monolayer expansion depending on the cell-substrate dissipation $D$ (initially 2500 cell system; stiffness parameters $\kappa_P = 0.12$, $\kappa_A = 0.18$; average cytoskeletal density $(Q+q)/2 = 35$; maximum cell polarity $\Delta Q = 10$; signalling radius $R = 2$; cytoskeletal update rate $\mu = 0.1$; cell-cell adhesion $B = 7$; cell-cell dissipation $\Delta B = 0$; cell-substrate adhesion penalty $\varphi = 0$; growth time $T_g = 180$; division time $T_d = 20$; size threshold for cell growth $A_T = 1\,A_{\text{ref}}$, where $A_{\text{ref}}$ is the cell size at equilibrium). **(A)** Maximal monolayer extension and roughness after an initial time of 200 MCS (it takes at least that long for first daughter cells to appear). **(B)** Time traces for selected values of $D$.

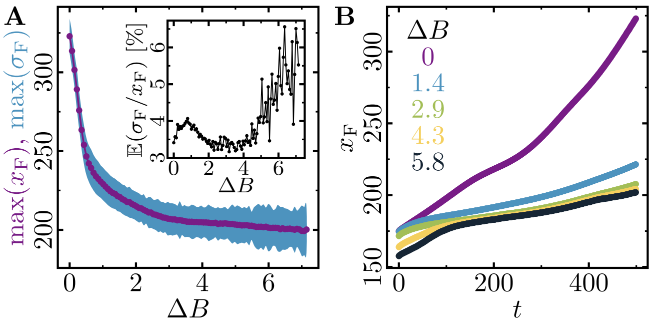


FIG. S14. Cell monolayer expansion depending on the cell-cell dissipation $\Delta B$ (initially 2500 cell system; stiffness parameters $\kappa_P = 0.12$, $\kappa_A = 0.18$; average cytoskeletal density $(Q+q)/2 = 35$; maximum cell polarity $\Delta Q = 10$; signalling radius $R = 2$; cytoskeletal update rate $\mu = 0.1$; cell-cell adhesion $B = 7$; cell-substrate dissipation $D = 0$; cell-substrate adhesion penalty $\varphi = 0$; growth time $T_g = 180$; division time $T_d = 20$; size threshold for cell growth $A_T = 1\,A_{\text{ref}}$, where $A_{\text{ref}}$ is the cell size at equilibrium). **(A)** Maximal monolayer extension and roughness after an initial time of 200 MCS (it takes at least that long for first daughter cells to appear). **Inset:** Relative roughness of the spreading monolayer relative to its size. **(B)** Time traces for selected values of $\Delta B$.

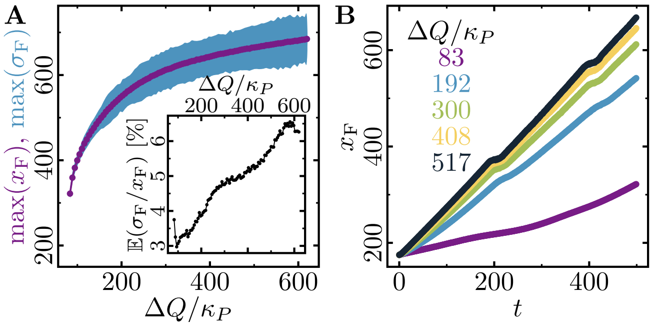


FIG. S16. Cell monolayer expansion depending on the maximum cell polarity $\Delta Q/\kappa_P$ (initially 2500 cell system; stiffness parameters $\kappa_P = 0.12$, $\kappa_A = 0.18$; average cytoskeletal density $(Q+q)/2 = 35$; signalling radius $R = 2$; cytoskeletal update rate $\mu = 0.1$; cell-cell adhesion $B = 7$; cell-cell dissipation $\Delta B = 0$; cell-substrate dissipation $D = 0$; cell-substrate adhesion penalty $\varphi = 0$; growth time $T_g = 180$; division time $T_d = 20$; size threshold for cell growth $A_T = 1\,A_{\text{ref}}$, where $A_{\text{ref}}$ is the cell size at equilibrium). **(A)** Maximal monolayer extension and roughness after an initial time of 200 MCS (it takes at least that long for first daughter cells to appear). **(B)** Time traces for selected values of $\Delta Q/\kappa_P$.

[1] F. Graner and J. A. Glazier, Phys. Rev. Lett. **69**, 2013 (1992).
[2] J. A. Glazier and F. Graner, Phys. Rev. E **47**, 2128 (1993).
[3] B. Alberts, A. Johnson, J. Lewis, M. Raff, K. Roberts, and P. Walter, *Molecular Biology of the Cell*, Molecular Biology of the Cell: Reference Edition No. Bd. 1 (Garland Science, 2008).
[4] D. Raucher and M. P. Sheetz, J. Cell Biol. **148**, 127 (2000).
[5] P. Friedl, Curr. Opin. Cell Biol. **16**, 14 (2004).
[6] N. B. Ouchi, J. A. Glazier, J.-P. Rieu, A. Upadhyaya, and Y. Sawada, Physica A **329**, 451 (2003).
[7] A. Mogilner, J. Math. Biol. **58**, 105 (2009).
[8] Currently, cell numbers up to $\mathcal{O}(10^4)$ can be achieved at acceptable computation times.
[9] A. F. M. Marée, A. Jilkine, A. Dawes, V. A. Grieneisen, and L. Edelstein-Keshet, Bull. Math. Biol. **68**, 1169 (2006).
[10] A. F. M. Marée, V. A. Grieneisen, and L. Edelstein-Keshet, PLoS Comput. Biol. **8**, 1 (2012).
[11] T. Pollard and G. Borisy, Cell **112**, 453 (2003).
[12] E. M. Kovacs, M. Goodwin, R. G. Ali, A. D. Paterson, and A. S. Yap, Current Biology **12**, 379 (2002).
[13] D. E. Leckband, Q. le Duc, N. Wang, and J. de Rooij, Curr. Opin. Cell Biol. **23**, 523 (2011).
[14] The energy difference associated with accepting $\mathcal{T}_{\rm rup}$ can be computed by standard means, simply using the substrate $\beta = -1$ as new target cell.
[15] J. F. Li and J. Lowengrub, Journal of Theoretical Biology **343**, 79 (2014).
[16] F. Barber, P.-Y. Ho, A. W. Murray, and A. Amir, Front. Cell. Dev. Biol. **5**, 92 (2017).
[17] B. I. Shraiman, Proc. Natl. Acad. Sci. U.S.A. **102**, 3318 (2005).
[18] J. Ranft, M. Basan, J. Elgeti, J.-F. Joanny, J. Prost, and F. Jülicher, Proc. Natl. Acad. Sci. U.S.A. **107**, 20863 (2010).
[19] N. Minc and M. Piel, Trends in Cell Biology **22**, 193 (2012).
[20] W. T. Gibson and M. C. Gibson, Curr. Top. Dev. Biol. **89**, 87 (2009).
[21] L. D. Landau, L. P. Pitaevskii, A. M. Kosevich, and E. Lifshitz, *Theory of Elasticity, Third Edition: Volume 7 (Course of Theoretical Physics)*, 3rd ed. (Butterworth-Heinemann, 1986).
[22] K. Keren, Z. Pincus, G. M. Allen, E. L. Barnhart, G. Marriott, A. Mogilner, and J. A. Theriot, Nature **453**, 475 (2008).
